# Binding of a blast fungus Zinc-finger fold effector to a hydrophobic pocket in the host exocyst subunit Exo70 modulates immune recognition in rice

**DOI:** 10.1101/2022.06.18.496527

**Authors:** Juan Carlos De la Concepcion, Koki Fujisaki, Adam R. Bentham, Neftaly Cruz Mireles, Victor Sanchez de Medina Hernandez, Motoki Shimizu, David M. Lawson, Sophien Kamoun, Ryohei Terauchi, Mark J. Banfield

## Abstract

Exocytosis plays an important role in plant-microbe interactions, both in pathogenesis and symbiosis. Exo70 proteins are integral components of the exocyst, an octameric complex that mediates tethering of vesicles to membranes in eukaryotes. Although plant Exo70s are known to be targeted by pathogen effectors, the underpinning molecular mechanisms and the impact of this interaction on infection is poorly understood. Here, we show the molecular basis of the association between the effector AVR-Pii of the blast fungus *Maganaporthe oryzae* and rice Exo70 alleles OsExo70F2 and OsExo70F3, which is sensed by the immune receptor pair Pii via an integrated RIN4/NOI domain. The crystal structure of AVR-Pii in complex with OsExo70F2 reveals that the effector binds to a conserved hydrophobic pocket in Exo70, defining a new effector/target binding interface. Structure-guided and random mutagenesis validates the importance of AVR-Pii residues at the Exo70 binding interface to sustain protein association and disease resistance in rice when challenged with fungal strains expressing effector mutants. Further, the structure of AVR-Pii defines a novel Zinc-finger effector fold (ZiF) distinct from the MAX fold previously described for the majority of characterized *M. oryzae* effectors. Our data suggests that blast fungus ZiF effectors bind a conserved Exo70 interface to manipulate plant exocytosis and that these effectors are also baited by plant immune receptors, pointing to new opportunities for engineering disease resistance.

**Significance statement:** Plant diseases destroy ∼20-30% of annual crop production, contributing to global food insecurity. Discovering how pathogen effectors target host proteins to promote virulence is essential for understanding pathogenesis and can be used for developing disease resistant crops. Here, we reveal the structural basis of how an effector from the blast pathogen (AVR-Pii) binds a specific host target (rice Exo70), and how this underpins immune recognition. This has implications for understanding the molecular mechanisms of blast disease and for the engineering of new recognition specificities in plant immune receptors to confer resistance to a major crop pathogen.

## Main text

Exocytosis is a cellular pathway in which membrane-bound vesicles are delivered from intracellular compartments to the plasma membrane for the release of their contents to the extracellular space (1). This pathway is essential for cell growth and division, as well as many other specialized processes that involve polarized secretion (1).

During exocytosis, a protein complex called the exocyst mediates the tethering of vesicles to the plasma membrane (2). The exocyst is an octamer formed by the proteins Sec3, Sec5, Sec6, Sec8, Sec10, Sec15, Exo70, and Exo84 (3), which assemble in a holocomplex (4–7) to control the spatiotemporal regulation of exocytosis, orchestrate cargo delivery and mediate vesicle secretion (8–11). The exocyst complex is conserved in all eukaryotes. In yeast and mammals, the exocyst component Exo70 is encoded by a single gene (12). However, plant Exo70s have expanded dramatically, resulting in high diversity in protein sequence and multiple gene copies (12, 13). This suggests that Exo70 proteins may have functionally diversified and adopted specialized functions in plants (10). Indeed, different Exo70s are involved in diverse plant processes including root development (14, 15), cell wall deposition (16, 17), symbiosis with arbuscular mycorrhiza (18), and cell trafficking pathways distinct from exocytosis such as autophagy (19–21).

Specific plant Exo70 proteins have been associated with disease resistance to pathogens and pests (13, 22–25). Like other components of cellular pathways involved in homeostasis and/or signalling, the exocyst complex is targeted by plant pathogens to promote disease and, in some cases, is actively monitored by the immune system. For example, Exo70 proteins can be guarded by plant receptors of the NLR (Nucleotide-Binding, Leucine-Rich repeat) superfamily (26–28) and have been shown to interact with RIN4, a well-known regulator of plant immunity (29, 30) that is also targeted by effectors from diverse pathogens (31, 32). Both Exo70 and RIN4 domains are also found as integrated domains in plant NLRs (13, 33–36), suggesting the importance of these two proteins in disease and plant defence.

The blast fungus pathogen *Magnaporthe oryzae* delivers effectors into the host to alter cellular processes, aiding successful colonization (37, 38). Genome sequencing uncovered hundreds of putative effectors harboured by this pathogen (39). However, only a small subset of these proteins have been functionally characterized to date. One such effector, AVR-Pii, interacts with two rice Exo70 subunits, OsExo70F2 and OsExo70F3, suggesting that the pathogen may target exocyst-mediated trafficking as a virulence-associated mechanism (28).

AVR-Pii was cloned alongside blast effectors AVR-Pik and AVR-Pia by association genetics (40). AVR1-CO39, AVR-Pik and AVR-Pia are founding members of the MAX (Magnaporthe Avrs and ToxB like) effector family (41), and studies on these effectors have been instrumental in defining the role of unconventional integrated domains in plant NLRs (42–49). Additional studies focused on these effectors and their cognate immune receptors have enabled engineering of bespoke immune responses to pathogen effectors (50–54). Despite the knowledge provided by structure/function studies of AVR-Pik and AVR-Pia, AVR-Pii has remained somewhat understudied.

Comprising only 70 residues, AVR-Pii is substantially smaller than AVR-Pik or AVR-Pia (40), and was not predicted to be a member of the MAX effector family (41). With remarkable specificity, AVR-Pii only associates with 2 alleles out of the 47 members of the rice Exo70 family (12), OsExo70F2 and OsExo70F3 (28). This suggests that AVR-Pii may target specific processes carried out by exocyst complexes harbouring these Exo70 alleles. Given that Exo70 proteins from phylogenetically distinct organisms share a common fold (55–57), AVR-Pii must exploit subtle structural differences to achieve this high interaction specificity. However, the molecular details of such stringent effector specificity are unknown. AVR-Pii is recognized by a rice disease resistance gene pair named Pii, which is comprised of the genetically linked genes Pii- 2 and Pii-1 (58). This recognition requires at least OsExo70F3 (28), and the association of AVR-Pii to OsExo70F3 is monitored by Pii through an unconventional RIN4/NOI domain integrated in the sensor NLR Pii-2 (59). However, the precise mechanism of recognition remains obscure.

Here, we focussed on elucidating the molecular basis of *M. oryzae* AVR-Pii interaction with rice Exo70 proteins. By determining the crystal structure of the effector in complex with OsExo70F2, we defined a new effector/target binding interface. We revealed that AVR-Pii adopts a Zinc-finger fold (ZiF) that has not been reported previously for plant pathogen effectors (60) and is distinct from the MAX fold found in the *M. oryzae* effectors whose structure is known to date (41). We then used structure-informed and random mutagenesis to dissect how the Exo70/AVR-Pii interface underpins effector binding, exploring the basis of effector specificity. Finally, we correlated Exo70/AVR-Pii binding with Pii-mediated resistance in rice, further strengthening the link between host virulence-associated targets and immune regulation. The exocyst complex is a target of diverse plant pathogen effectors and is linked to resistance against pathogens and pests, suggesting a role as a “hub” in pathogenesis and plant defence. Our study expands our understanding of the molecular mechanisms used by plant pathogen effectors to target host proteins and may enable new approaches to engineering of disease resistance.

## Results

### AVR-Pii binds OsExo70F2 and OsExo70F3 with high affinity and specificity

To explore a detailed understanding of the interaction between AVR-Pii and rice Exo70s, we performed a Yeast-2-Hybrid assay (Y2H) co-expressing the effector with the rice Exo70 alleles OsExo70B1, OsExo70F2 and OsExo70F3. Consistent with previous work (28), we observed yeast growth and the development of blue colouration with X-α-gal, both readouts of protein-protein interactions, for AVR-Pii co-expressed with OsExo70F2 or OsExo70F3, but not with OsExo70B1 **(Figure 1a)**.

**Figure 1.**
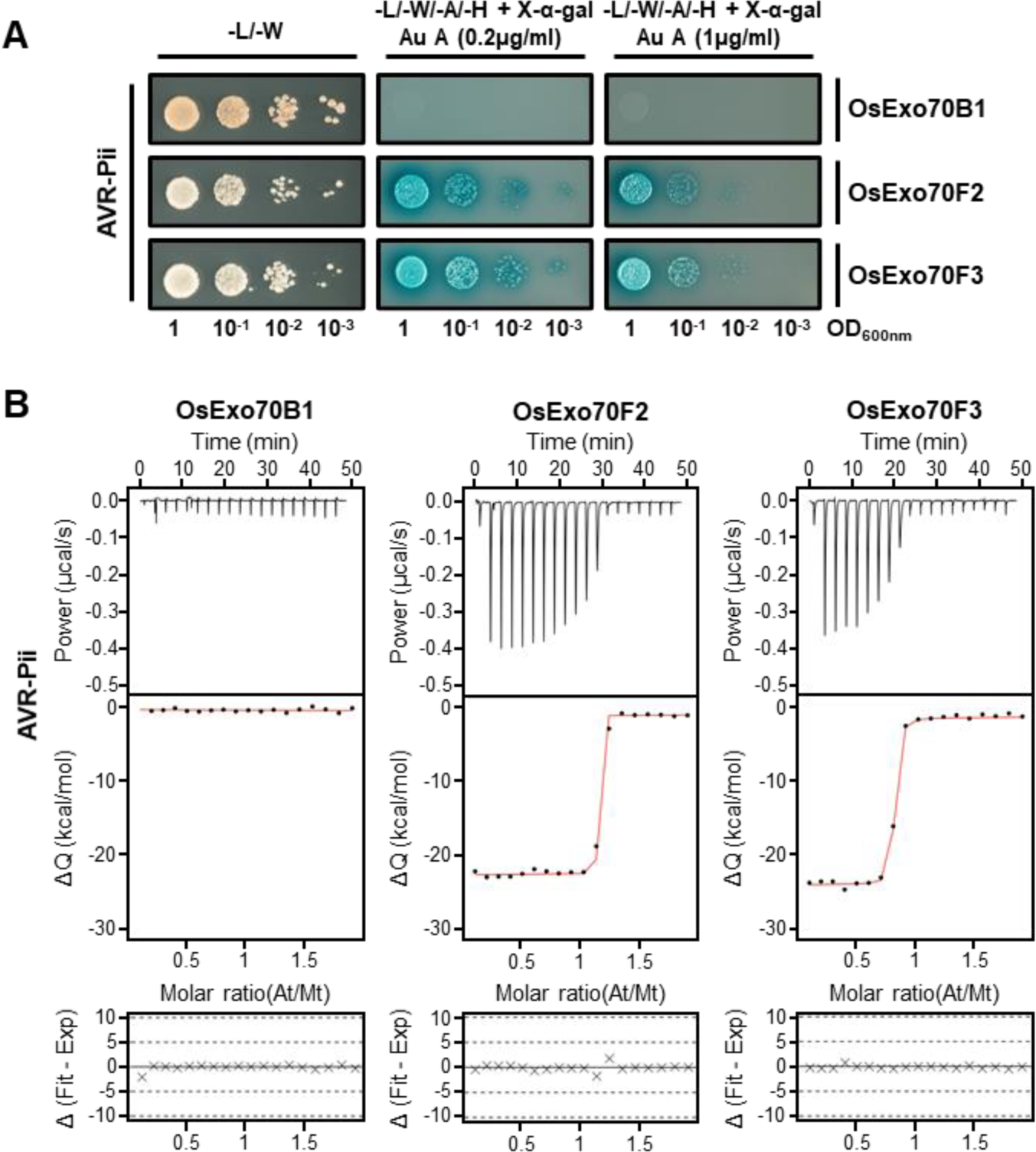
AVR-Pii binds specifically to rice Exo70F2 and Exo70F3 in yeast and in vitro. **(A)** Yeast-Two-Hybrid assay of AVR-Pii with rice OsExo70B1, OsExo70F2 and OsExo70F3. The control plate for yeast growth is on the left, with quadruple dropout media supplemented with X-α-gal and increasing concentrations of Aureobasidine A (Au A) on the right. Growth and development of blue colouration in the selection plate are both indicative of protein:protein interactions. OsExo70 proteins were fused to the GAL4 DNA binding domain, and AVR-Pii to the GAL4 activator domain. Each experiment was repeated a minimum of three times, with similar results. **(B)** Binding of AVR-Pii to OsExo70 proteins determined by ITC. Upper panels show heat differences upon injection of AVR-Pii into the cell containing the respective OsExo70 allele. Middle panels show integrated heats of injection (dots) and the best fit (solid line) using to a single site binding model calculated using AFFINImeter ITC analysis software (79). Bottom panels represent the difference between the fit to a single site binding model and the experimental data; the closer to zero indicates stronger agreement between the data and the fit. The experiments shown are representative of three replicates. Other replicates for each experiment are shown in **Figure S5**. The thermodynamic parameters obtained in each experiment are presented in **Table S1**.

This confirms that AVR-Pii specifically interacts with these Exo70 alleles. Growth of yeast was also clearly observed at elevated concentrations of Aureobasidin A **(Figure 1a)**, suggesting that the association of AVR-Pii with OsExo70F2 and OsExo70F3 is robust.

To test for effector/target interactions in vitro, we optimised a pipeline to produce and purify rice Exo70 subunits OsExo70B1, OsExo70F2 and OsExo70F3 by heterologous expression in *E. coli* **(Figure S1)** (see details in **SI materials and methods**). Analytical gel filtration analysis of purified rice Exo70 alleles showed that proteins with truncated N-terminal domains elute as a monodisperse peaks, suggesting they are suitable for further biophysical experiments **(Figure S2)**. Likewise, we purified the effector domain of AVR-Pii (residues 20 to 70) adapting a protocol previously used for the purification of the blast effector AVR-Pik **(Figure S3)** (see details in **SI materials and methods**). The molecular mass of the effector was confirmed by intact mass spectrometry **(Figure S4)**.

To investigate the strength of binding between AVR-Pii and Exo70 alleles we used Isothermal Titration Calorimetry (ITC). We measured heat differences (indicative of protein/protein interactions) after titration of the AVR-Pii effector into a solution containing purified rice Exo70 proteins and used this information to calculate K*_d_* values for the interaction. These experiments showed AVR-Pii binds to both OsExo70F2 and OsExo70F3 with nanomolar affinity **(Figure 1b, Figure S5, Table S1)**.

No interaction was detected between AVR-Pii and OsExo70B1 **(Figure 1b, Figure S5, Table S1)**, confirming the high specificity of the binding observed in Y2H.

In summary, we confirmed that AVR-Pii is a selective effector that binds to a specific subset of allelic rice Exo70s with high affinity.

### Crystal structure of AVR-Pii in complex with OsExo70F2

After confirming that AVR-Pii binds to OsExo70F2 and OsExo70F3 in vitro, we showed that an OsExo70/AVR-Pii complex can be reconstituted and purified to homogeneity **(Figure S6)**. A reconstituted OsExo70F2/AVR-Pii complex was stable and could reach high concentrations. Using this sample, we obtained protein crystals that diffracted X-rays to 2.7 Å resolution at the Diamond Light Source (DLS, Oxford, UK). Details of the X-ray data collection, structure solution and structure completion are given in the **SI materials and methods** section and **Table S2**.

In the crystal structure, AVR-Pii and OsExo70F2 form a 1:1 complex **(Figure 2a)**. OsExo70F2 adopts an elongated rod-like shape formed by the stacking of four domains (Domains A – D), with 16 α-helices in total (annotated α1 – α16) arranged in four-helix bundles **(Figure S7a)**. Despite significant differences in sequence, the OsExo70F2 structure closely resembles the fold of the *Arabidopsis* AtExo70A1 protein (PDB ID: 4L5R) (57); and the mouse MmExo70 protein (PDB ID: 2PFT) (56), which can be aligned to OsExo70F2 structure with an R.M.S.D. of 1.13 Å and 1.00 Å across 188 and 177 pruned atom pairs respectively as calculated with ChimeraX (61) **(Figure** S7b).

**Figure 2.**
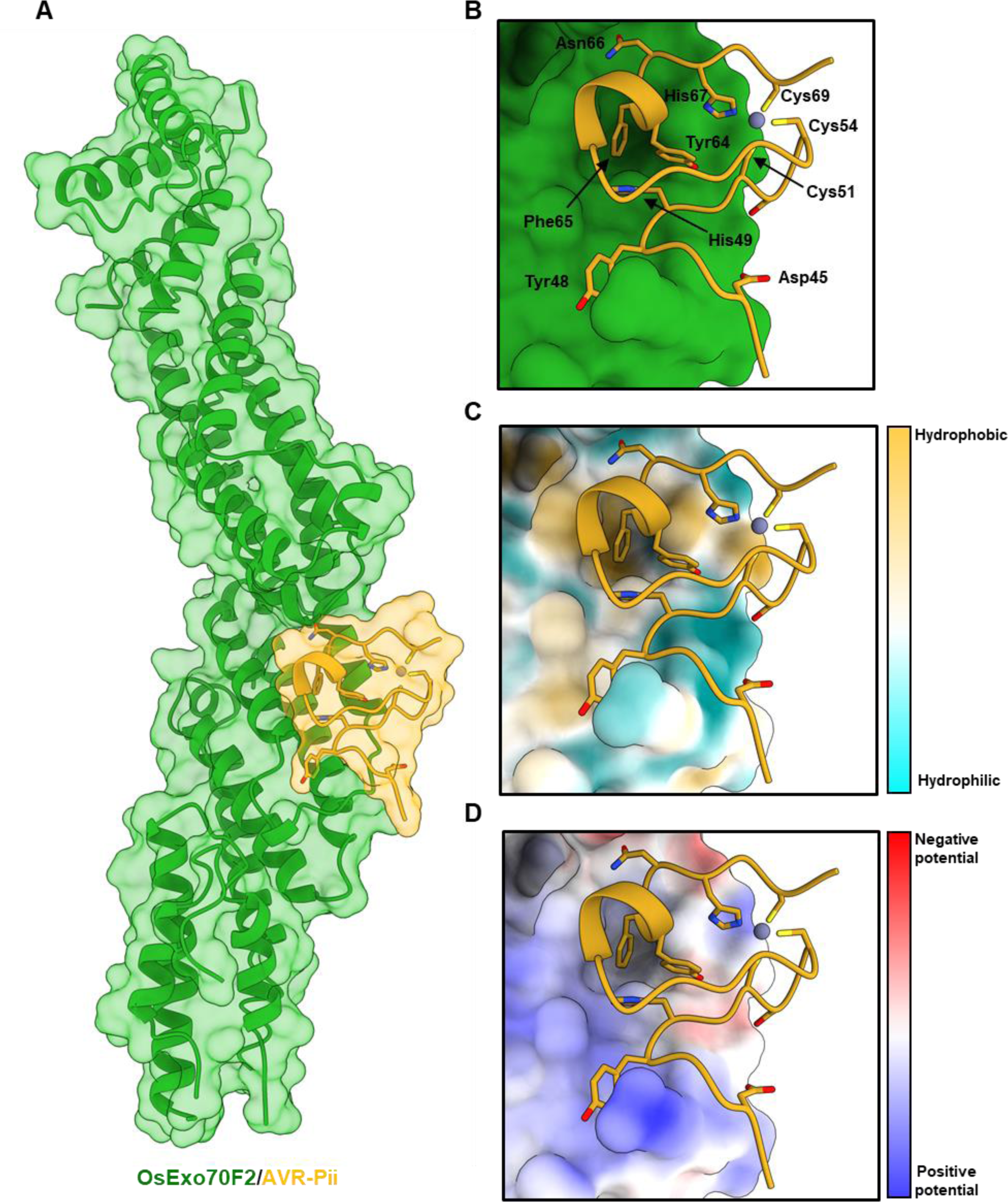
The crystal structure of OsExo70F2 in complex with AVR-Pii reveals hydrophobic residues dominate the interaction interface. (A) Schematic representation of OsExo70F2 in complex with AVR-Pii. Both molecules are represented as cartoon ribbons, with the molecular surface also shown and coloured as labelled in green and yellow for OsExo70F2 and AVR-Pii, respectively. **(B)** Close-up view of the interaction interface between OsExo70F2 and AVR-Pii. OsExo70F2 is presented as a solid surface, with the effector as cartoon ribbons and side chains displayed as a cylinder for AVR-Pii interacting residues (Asp45, Tyr48, His49, Tyr64, Phe65 and Asn66) in additional to the residues co-ordinating the Zn^2+^ atom (Cys51, Cys54, His67 and Cys69). **(C)** OsExo70F2 surface hydrophobicity representation at the AVR-Pii interaction interface, residues are coloured depending on their hydrophobicity from light blue (low) to yellow (high). **(D)** Representation of OsExo70F2 surface electrostatic potential at the AVR-Pii interaction interface, residues are coloured depending on their electrostatic potential from dark blue (positive) to red (negative).

Consistent with existing Exo70 structures, the N-terminus of OsExo70F2 is disordered, and the first residue modelled in the electron density is Ser85. Further, several additional loop regions were disordered, including those connecting the helices α2 and α3 (residues 130 to 156), α3 and α4 (231 to 256), α6 and α7 (330 and 331), α10 and α11 (461 to 482), α13 and α14 (572 to 598), and α15 and α16 (648 to 659).

Following placement of OsExo70F2, we identified electron density consistent with the sequence of AVR-Pii and were able to build residues 44 to 70 of the effector (residues 20 to 43 were not observed in the electron density). This C-terminal region of AVR-Pii revealed a novel fold for a *M. oryzae* effector based on a Zinc-finger motif **(Figure 2, Figure S8)**. This fold is sustained by AVR-Pii residues Cys51, Cys54, His67 and Cys69 that coordinate a Zn^2+^ atom **(Figure 2b, Figure S8)**. A structural similarity search performed with PDBeFold (62) revealed the AVR-Pii structure is most similar to LIM domain Zinc-fingers with a motif of C-X_2_-C-X_12_-H-X-C, however, it lacks a second zinc binding motif commonly found in this class of domains. We refer to the AVR-Pii fold as ZiF for Zinc-finger Fold and note that this 3D structure has not been previously reported for other plant pathogen effectors (60), and is distinct from the MAX fold found for other *M. oryzae* effectors whose structures are known (41).

### AVR-Pii interacts with OsExo70F2 via a hydrophobic pocket

The binding interface between OsExo70F2 and AVR-Pii is well resolved in the electron density. AVR-Pii locates to an amphipathic surface formed at the junction of OsExo70F2 domains B and C **(Figure 2, Figure S8)**, with residues from helices α8, α9, and α10 contributing to the effector-binding interface **(Figure 2b, Figure S8)**. Analysis of the complex using QtPISA (63) reveals both hydrophobic and hydrogen bond interactions in the complex, with a remarkable 20 of 27 AVR-Pii residues (74%) involved in contacts with OsExo70F2 (**Figure S9, Figure S10**). Further, molecular lipophilicity potential and Coulombic electrostatic potential (calculated with ChimeraX (61)) reveals distinct hydrophobic and charged regions on the surface of OsExo70F2 surrounding the AVR-Pii binding interface **(Figure 2c, d)**. The most striking feature of the OsExo70F2/AVR-Pii interface is a hydrophobic pocket formed by the OsExo70F2 residues Phe416, Leu420, Met437, Tyr440 and Val441, which accommodates AVR-Pii Phe65, and to a lesser extent, Tyr64 **(Figure 2b, c and d)**.

### Amino acid variation in the OsExo70 hydrophobic pocket underpins AVR-Pii binding specificity

To better understand how AVR-Pii achieves a high binding specificity towards different Exo70 alleles, we analysed conservation of residues at the OsExo70/AVR-Pii binding interface using Consurf (64). Unexpectedly, this analysis showed significant conservation in the residues at the AVR-Pii interface, with the residues surrounding the hydrophobic pocket showing limited variability **(Figure S11)**. We then modelled rice OsExo70B1 using AlphaFold2 (65), as implemented in ColabFold (66) **(Figure S12),** to observe whether structural homology could help understand specificity. While the N-terminal region of OsExo70B1 could not be accurately resolved,

AlphaFold2 produced a high confidence model for the domains present in the OsExo70F2/AVR-Pii complex, including the effector binding interface. Side-by-side comparison of the sequence and structure of OsExo70F2 with the model of OsExo70B1 showed small differences in the residues forming the hydrophobic pocket, with OsExo70F2 Phe416, Val419, Leu420 and Met437 replaced by Leu405, Leu408, Ile 409 and Ile426 at equivalent positions in OsExo70B1 **(Figure S13a)**. While these polymorphisms do not appear to alter the overall hydrophobicity or electrostatic potential at the effector binding interface **(Figure S13)**, the hydrophobic pocket of OsExo70F2 is not observed in the OsExo70B1 model **(Figure S13)**. This suggests that AVR-Pii residues Tyr64 and Phe65 could not be accommodated, resulting in the lack of binding to OsExo70B1. Therefore, we conclude that AVR-Pii specificity is underpinned by small changes in the Exo70 binding interface that dramatically alter binding affinity.

### Mutations at the Exo70/AVR-Pii interface prevent binding

Prior to obtaining the structure of the OsExo70F2/AVR-Pii complex we used random mutagenesis coupled with Y2H to identify AVR-Pii residues involved in binding to OsExo70 proteins **(Figure S14)**. Using this approach, we obtained five independent AVR-Pii mutants within the effector domain, named M1 to M5 **(Figure S14a)**. These mutants showed reduced (M2) or abrogated (M4 and M5) binding to OsExo70F3 **(Figure S14b)**. Western blot analysis showed that M2 and M5 displayed a lower protein accumulation in yeast **(Figure S14c).** As M2 and M5 carry mutations in Zn^2+^ binding residues (Cys54Arg in M2 and Cys51Arg in M5), it is likely these affect protein folding and protein stability. AVR-Pii M4 was the only mutant displaying lack of binding without compromised effector accumulation. This mutant harbours two residue changes, Arg43Ser and Tyr64Asp. We therefore generated single mutants Arg43Ser, Arg43Ala, Tyr64Asp and Tyr64Ala to investigate the contribution of these residues to the binding to OsExo70F3. Y2H assays show that only Tyr64Asp prevented AVR-Pii binding to OsExo70F3 **(Figure S14d)**, and none of these mutations affected protein accumulation **(Figure S14e)**.

Then, based on the crystal structure we designed point mutants in AVR-Pii residues Tyr64 (Tyr64Arg) and Phe65 (Phe65Glu) as these were the dominant residues revealed at the interface. Y2H assays showed that AVR-Pii Tyr64Arg and Phe65Glu mutations severely affected binding to OsExo70F2 and prevented binding to OsExo70F3 **(Figure 3a)**. These mutations did not affect protein accumulation of the effector in yeast cells **(Figure S15)**. To extend the Y2H analysis, we expressed and purified AVR-Pii Tyr64Arg and Phe65Glu mutants and tested their ability to bind OsExo70F2 and OsExo70F3 by ITC. Consistent with the Y2H assays, Tyr64Arg and Phe65Glu mutations impacted binding of AVR-Pii to OsExo70F2 and OsExo70F3, with essentially no binding observed using this technique **(Figure 3b, Figure S16, Figure S17 and Table S3)**. Together, these experiments confirmed the AVR-Pii residues that locate to the OsExo70F2 hydrophobic pocket are essential for target binding and effector specificity.

**Figure 3.**
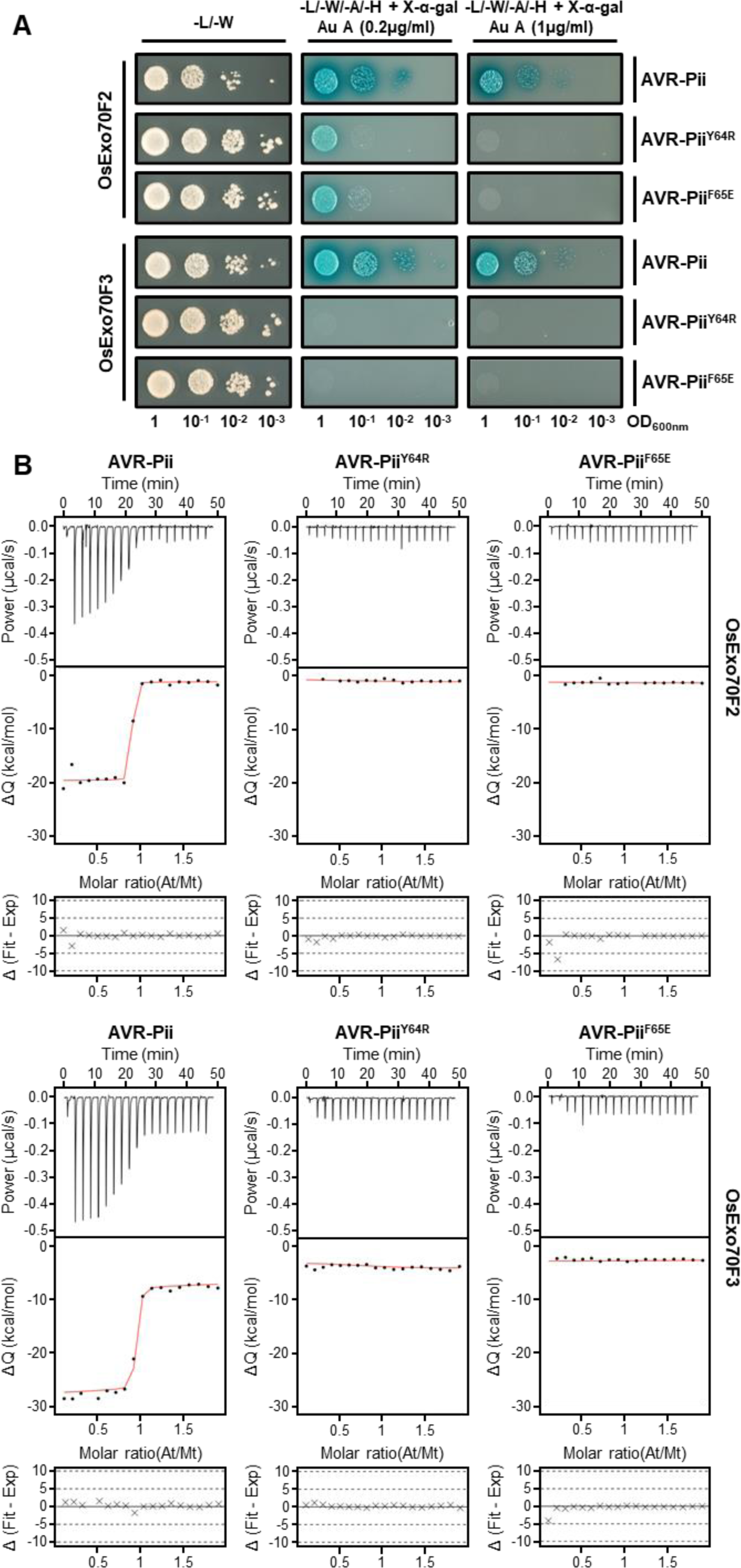
Mutations at the AVR-Pii binding interface perturb interactions with rice Exo70 proteins. **(A)** Yeast-Two-Hybrid assay of AVR-Pii mutants Tyr64Arg and Phe65Glu with rice OsExo70F2 and OsExo70F3. The control plate for yeast growth is on the left, with quadruple dropout media supplemented with X-α-gal and increasing concentrations of Aureobasidine A (Au A) on the right. Growth and development of blue colouration in the selection plate are both indicative of protein:protein interactions. Wild-type AVR-Pii is included as positive control. Exo70 proteins were fused to the GAL4 DNA binding domain, and AVR-Pii to the GAL4 activator domain. Each experiment was repeated a minimum of three times, with similar results. **(B)** Binding of AVR-Pii mutants Tyr64Arg and Phe65Glu to rice OsExo70F2 and OsExo70F3 determined by isothermal titration calorimetry (ITC). Wild type AVR-Pii was included as positive control. Upper panels show heat differences upon injection of AVR-Pii mutants into the cell containing the respective OsExo70 allele. Middle panels show integrated heats of injection (dots) and the best fit (solid line) using to a single site binding model calculated using AFFINImeter ITC analysis software (79). Bottom panels represent the difference between the fit to a single site binding model and the experimental data; the closer to zero indicates stronger agreement between the data and the fit. Panels are representative of three replicates. Other replicates for each experiment are shown in **Figure S16 and S17**. The thermodynamic parameters obtained in each experiment are presented in **Table S3**.

### Mutations at the Exo70/AVR-Pii interface abrogate Pii-mediated resistance to rice blast

Having identified residues that prevent Exo70/AVR-Pii binding by Y2H and in vitro, we transformed the *M. oryzae* Sasa2 strain (which lacks AVR-Pii) with wild-type AVR-Pii and the Tyr64Arg and Phe65Glu mutants to observe the impact on resistance mediated by the Pii NLR pair. For this assay, different independent Sasa2 transformants were recovered and their virulence tested in the susceptible rice cultivar Moukoto. We discarded non-infective transformants and performed RT-PCR to test for expression of the effector in the remaining strains **(Figure S18)**.

Infective strains expressing AVR-Pii wild-type, Tyr64Arg or Phe65Glu were then spot inoculated on rice cultivars Moukoto (lacking Pii resistance) and Hitomebore (harbouring Pii resistance). The length of lesions was measured 10 days post infection to assay the extent of disease progression **(Figure 4)**. As expected, all strains were virulent on the susceptible rice cultivar Moukoto, but the strains expressing wild-type AVR-Pii did not form expanded lesions on Hitomebore. However, we observed that AVR-Pii mutants that abrogate binding to OsExo70F2 and OsExo70F3 in Y2H and in vitro assays are not recognized in Hitomebore, with large disease lesions forming equivalent in size to those observed for untransformed Sasa2 **(Figure 4, Figure S19, Figure S20, Figure S21)**. This data corroborates the direct link between OsExo70/AVR-Pii binding and recognition by the rice NLR pair Pii.

**Figure 4.**
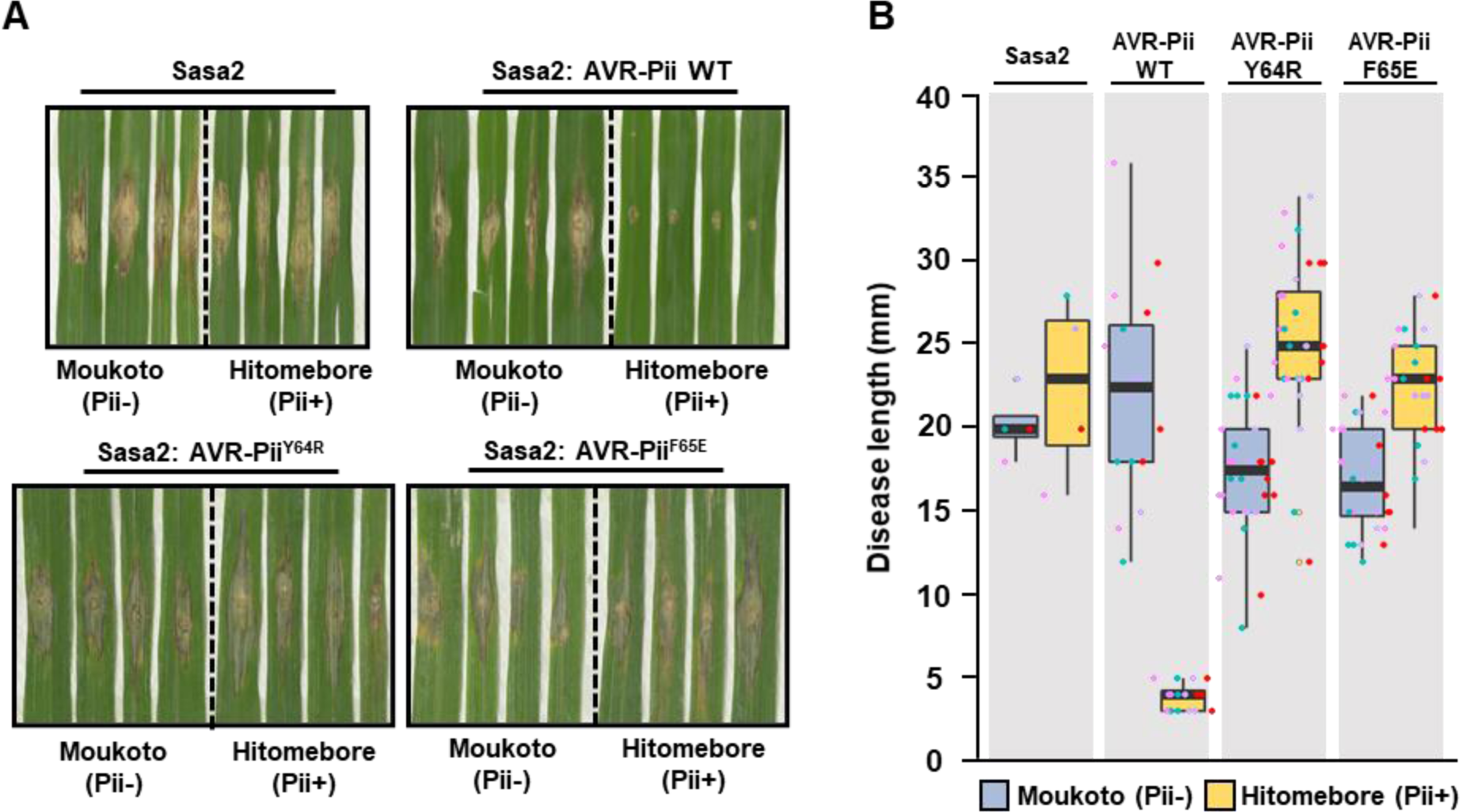
Mutations at the binding interface of AVR-Pii with OsExo70 abrogate recognition by Pii resistance in rice. (A) Rice leaf blade spot inoculation of transgenic *M. oryzae* Sasa2 isolates expressing AVR-Pii, AVR-Pii Tyr64Arg or AVR-Pii Phe65Glu in rice cultivars Moukoto (Pii-) and Hitomebore (Pii+). For each experiment, representative images from replicates with independent *M. oryzae* transformants are shown. Wild-type *M. oryzae* isolate Sasa2 is included as a control. Images for each replicate of AVR-Pii, AVR-Pii Tyr64Arg and AVR-Pii Phe65Glu are presented in **Figure S19-S21**. **(B)** Measurement of vertical length of the disease lesion caused by *M. oryzae* Sasa2 as well as transgenic *M. oryzae* Sasa2 isolates harbouring AVR-Pii, AVR-Pii Tyr64Arg or AVR-Pii Phe65Glu in rice cultivars Moukoto (Pii-) and Hitomebore (Pii+). Lesions in rice cultivars Moukoto (Pii-) and Hitomebore (Pii+) are represented by blue and yellow boxes, respectively. For each isolate, a total of four biological replicates were performed, and the data are presented as box plots. The centre line represents the median, the box limits are the upper and lower quartiles, the whiskers extend to the largest value within Q1 - 1.5× the interquartile range (IQR) and the smallest value within Q3 + 1.5× IQR. All the data points are represented as dots with distinct colours for each biological replicate.

## Discussion

Exocytosis has recently emerged as a conserved eukaryotic trafficking pathway with roles in plant immunity and symbiosis (18, 21–24). The exocyst complex is targeted by effectors from diverse pathogens (28, 67), and Exo70 domains have been integrated in plant NLRs (33, 34), likely to act as effector baits for the detection of pathogens by the immune system. Therefore, understanding the molecular and structural basis of manipulation of plant exocytosis by pathogen effectors has the potential to uncover new mechanisms of pathogen virulence and, ultimately, may pave new ways for engineering the outcome of plant-pathogen interactions (50, 52, 54, 68). In this study, we uncovered the molecular details of how the blast pathogen effector AVR-Pii binds to rice exocyst subunit Exo70 and we dissected the structural determinants of this interaction.

Although hundreds of putative effectors are encoded in pathogen genomes, only limited examples of plant pathogen effector-host target interactions have been dissected in molecular detail to date (69). Intriguingly, some pathogen effectors have evolved to target specific members of large host protein families, potentially to avoid compromising host cell viability. AVR-Pii is an example of an effector with a striking target specificity, associating with only 2 out of 47 members of the rice Exo70 protein family (Fujisaki et al., 2015). By obtaining the structure of the OsExo70F2/AVR-Pii complex we revealed the molecular basis of such a high specificity. Surprisingly, the effector binds to a moderately conserved region of the Exo70 domain, but one that contains subtle differences between alleles. Therefore, AVR-Pii appears to have evolved high specificity by exploiting small differences at the Exo70 binding interface, specifically within a hydrophobic pocket that allows for the docking of the effector.

Although a precise function of the exocyst complex in plant immunity remains yet to be described, some Exo70 alleles have been shown to associate with RIN4, a well- known target of multiple pathogen effectors that is guarded by the plant immune system (29, 30). An increasing number of effectors have been reported to alter the Exo70-RIN4/NOI immune node (30) and, much like RIN4, some Exo70s activate the plant immune system upon perturbation of their function (26).

The structure of AVR-Pii reported here reveals a new protein fold for fungal effectors, based on a Zinc Finger domain, that differs from the MAX fold shared by all the *M. oryzae* effectors whose structure is known to date (41, 43, 70, 71). Zinc Finger domains are abundant in nature, and can be regularly found as single domains in larger multi-domain proteins that are implicated in a variety of processes, from DNA interaction to signalling hubs and protein-protein scaffolds that regulate cellular functions, such as autophagy or G protein-coupled receptor signalling (72, 73). While the AvrP effector from Flax rust, *Melampsora lini*, also has been reported to have a Zinc finger fold (74), these two proteins share no structural similarity.

AVR-Pii targets OsExo70F2 and OsExo70F3, suggesting the effector may interfere with a function related to a unique role of these Exo70 alleles. However, any specific function of OsExo70F2 or OsExo70F3, compared with other rice Exo70 alleles remains to be determined. Structural modelling based on the Cryo-EM model of the yeast exocyst (4) suggests that AVR-Pii would sit on the outer side of the complex, where it would not be expected to disrupt the association of the subunits forming subcomplex I (Sec3, Sec5, Sec6 and Sec8) or subcomplex II (Sec10, Sec15, Exo70 and Exo84) **(Figure S22a)**. However, AVR-Pii is found close to the interface between Sec5 and Exo70 which may interfere with the necessary assembly of both exocyst subcomplexes into the holocomplex for the late stages of exocytosis (11) **(Figure S22b)**. This suggests a function by which AVR-Pii may target assembly of exocyst complexes equipped with Exo70 subunits OsExo70F2 and OsExo70F3, but further experiments are required to investigate this.

Plant immune receptors have been the focus of engineering to improve disease resistance to some of the most destructive pathogens of crops (75), with limited success. Recent studies of *M. oryzae* MAX effectors, particularly AVR-Pik and AVR- Pia that target HMA domain containing proteins (69, 76, 77), and are bound by integrated HMA domains in NLR receptors (44, 46, 47), have demonstrated proof-of-concept for engineering NLRs to generate expanded recognition profiles (50–54, 78). Like HMA domains, Exo70s are found as integrated domains in some plant NLRs (33, 34). In addition to defining a new effector fold and determining the structural basis of a new effector/target interface, our results will help uncover new approaches for NLR engineering, for example, by repurposing integrated Exo70 domains to perceive different effectors.

## Materials and methods

### Gene cloning

Detailed information for gene cloning is provided in the **SI Materials and Methods**.

### Protein expression and purification

A full protocol for the heterologous expression and purification of rice Exo70 proteins and *M. oryzae* AVR-Pii effector is provided in the **SI Materials and Methods** section.

### Crystallization, data collection and structure solution

Details for X-ray data collection, structure solution and structure completion are given in the **SI Materials and Methods**.

### Protein-protein interaction: Yeast-2-hybrid

To detect protein–protein interactions between rice Exo70 proteins and AVR-Pii effectors in a yeast two-hybrid system, we used the Matchmaker Gold System (Takara Bio USA) following a protocol adapted from (46) and are detailed in **SI Materials and**

## Methods

### Protein-protein interaction: Isothermal titration calorimetry

ITC experiments were performed using a MicroCal PEAQ-ITC (Malvern, UK). In each case 300 μl of OsExo70 at 10 μM were placed in the calorimetric cell and titrated with 100 μM of AVR-Pii wild-type or mutants in the syringe. Each run included a single injection of 0.5 μL followed by 18 injections of 2 μL each at intervals of 120 second with a stirring speed of 750 rpm. Data were processed with AFFINImeter ITC analysis software (79). ITC runs for wild-type and mutants were done in triplicate at 25°C in 20 mM HEPES (pH 7.5), 150 mM NaCl and 5% (vol/vol) glycerol buffer supplemented with 1 mM TCEP.

### Rice blast infection assay

Conidial suspension (2-5 × 10^5^ conidia/ml) were prepared from the transgenic *M. oryzae* and used for leaf blade spot inoculation using rice cultivar Hitomebore (Pii+) and Moukoto (Pii-) as described previously (80). Disease lesions were photographed 10 days after inoculation, and vertical length was measured.

### RT-PCR

For RT-PCR analysis, the total RNA was extracted from the disease lesion-containing Moukoto leaves, and cDNA were synthesized using oligo dT primer. RT-PCR was performed using a specific primer set for AVR-Pii, and for *M. oryzae* Actin as control.

## Acknowledgments

We thank the Diamond Light Source, UK (beamline i03 under proposal MX18565) for access to X-ray data collection facilities. Dr Clare Stevenson (JIC X-ray Crystallography/Biophysical Analysis Platform) for help with X-ray data collection and ITC and Dr Gerhard Saalbach (JIC Proteomics facility) for intact mass analysis of AVR-Pii. We also thank Dr. Indira Saado for critical reading of the manuscript. This work was supported by the UKRI Biotechnology and Biological Sciences Research Council (BBSRC, UK, grants BB/P012574, BBS/E/J/000PR9795, BBS/E/J/000PR9777, BB/V015508/1), the European Research Council (proposal 743165), the John Innes Foundation, the Gatsby Charitable Foundation, the European Commission through the Erasmus+ programme, and JSPS Grant 20H05681.

## Data availability

Protein structure of rice blast effector protein AVR-Pii in complex with the host target Exo70F2 from rice, and the data used to derive this, have been deposited at the Protein DataBank (PDB) with accession code 7PP2.

## Supplementary information

### Extended materials and methods

#### Gene cloning

For protein production in *E. coli,* codon optimised AVR-Pii (amino acid residues Leu20 to Asn70) was synthesised (GenScript) and subsequently cloned into the pOPINM vector (81) using the In-Fusion cloning kit (Takara Bio USA). AVR-Pii Trp64Arg and Phe65Glu were synthesised as PCR products (Gblocks, IDT) and cloned into pOPINM in the same way. Truncated versions of rice Exo70 alleles OsExo70B1^Δ91^, OsExo70F2^Δ83^ and OsExo70F3^Δ93^ were generated using standard molecular biology techniques from appropriate templates described by Fujisaki et al. (28) and cloned into pOPINS3C (81) using the In-Fusion cloning kit (Takara Bio USA).

For Y2H, wild-type and mutant AVR-Pii effectors (amino acid residues Leu20 to Asn70) were cloned in pGADT7 while full length CDS of rice Exo70 alleles OsExo70B1, OsExo70F2 and OsExo70F3 were cloned in pGBKT7. In both cases, plasmids were linearized by double digestion with EcoRI and BamHI (New England Biolabs) and genes of interest were introduced using the In-Fusion cloning kit (Takara Bio USA).

For random mutagenesis we used the Diversify PCR Random Mutagenesis Kit (Takara Bio USA) and subsequently cloned the mutagenized AVR-Pii PCR fragments in pGADT7 as described above.

To construct pCB1531 plasmids with AVR-Pii wild-type, Trp64Arg and Phe65Glu under control of pex22 promotor (40), coding sequences were amplified from appropriate templates by PCR using primer set KF1341f (5’GCTCTAGAAAAATGCAACTTTCCAAAATTAC3’) and KF1295r (5’CGGGATCCTTAGTTGCATTTATGATT 3’). The PCR products were digested by BamHI and XbaI and inserted into the vector pCB1531-pex22p-EGFP (40) linearized with BamHI and XbaI. Resulting plasmids (pCB1531-pex22p-AVR-Pii-WT, Y64R and F65E) were transformed into *M. oryzae* Sasa2 strain lacking AVR-Pii gene as described previously (82).

#### Expression and purification of proteins for X-ray crystallography and in vitro binding studies

To enable the study of the OsExo70/AVR-Pii interactions in vitro, we produced stable rice OsExo70 proteins in *E. coli*. SUMO-tagged OsExo70 alleles with the predicted N-terminal α-helix truncated encoded in pOPINS3C were produced in *E. coli* Rosetta™ (DE3). Cell cultures were grown in autoinduction media (83) at 37°C for 5–7 hr and then at 16°C overnight. Cells were harvested by centrifugation and re-suspended in 50 mM Tris-HCl (pH 7.5), 500 mM NaCl, 50 mM glycine, 5% (vol/vol) glycerol, and 20 mM imidazole supplemented with EDTA-free protease inhibitor tablets (Roche). Cells were sonicated and, following centrifugation at 40,000xg for 30 min, the clarified lysate was applied to a HisTrap™ Ni2+-NTA column connected to an AKTA Xpress purification system (GE Healthcare). Proteins were step-eluted with elution buffer (50 mM Tris-HCl (pH7.5), 500 mM NaCl, 50 mM glycine, 5% (vol/vol) glycerol, and 500 mM imidazole) and directly injected onto a Superdex 200 26/60 gel filtration column pre-equilibrated with 20 mM HEPES (pH 7.5), 150 mM NaCl and 5% (vol/vol) glycerol supplemented with 1mM TCEP. Elution fractions were collected and evaluated by SDS-PAGE, revealing a band close to 70 kDa **(Figure S1a)**. Fractions were combined and incubated overnight with 3C protease (10 μg/mg fusion protein).

Rice Exo70 proteins were separated from the SUMO tag by passing the protein mixture solution through a HisTrap™ Ni2+-NTA column equilibrated with 50 mM Tris-HCl (pH 7.5), 500 mM NaCl, 50 mM glycine, 5% (vol/vol) glycerol, and 20 mM imidazole **(Figure S1b)**. Exo70 proteins were mainly present in the first and second wash-through **(Figure S1b)** from the column, whilst the SUMO tag was retained until the final elution with elution buffer **(Figure S1b)**. Fractions containing Exo70 proteins were pooled together and concentrated for further purification and buffer exchange by gel filtration onto a Superdex 200 16/60 column pre-equilibrated with 20 mM HEPES (pH 7.5), 150 mM NaCl and 5% (vol/vol) glycerol supplemented with 1mM TCEP **(Figure S1c)**. Fractions containing purified Exo70 proteins were combined and concentrated for structural and biophysical studies.

For OsExo70 gel filtration analysis, a volume of 110 μl of each sample was separated at 4 °C on a Superdex 200 10/300 size exclusion column (GE Healthcare), pre- equilibrated in 20 mM HEPES (pH 7.5), 150 mM NaCl, 1 mM TCEP and 5% (vol/vol) glycerol and at a flow rate of 0.5 ml/min. Fractions of 0.5 ml were collected for analysis by SDS–PAGE.

MBP-tagged effector domain (Amino acid residues 20 to 70) for wild-type AVR-Pii, Trp64Arg and Phe65Glu encoded by the pOPINM constructs were produced in *E. coli* SHuffle cells (84). Cell cultures were grown in autoinduction media (83) at 30°C for 5–7 hr and then at 16°C overnight. After harvest by centrifugation, cells were resuspended and disrupted as described above for OsExo70 expression.

The soluble fusion protein 6xHis:MBP:AVR-Pii was be purified from *E. coli* cell lysates by IMAC on a HisTrap™ Ni2+-NTA column connected to an AKTA Xpress purification system (GE Healthcare) coupled with gel filtration onto a Superdex 75 26/60 gel filtration column pre-equilibrated with 20 mM HEPES (pH 7.5), 150 mM NaCl and 5% (vol/vol) glycerol **(Figure S3a)**. The fractions containing the eluted protein were subsequently treated with 3C protease as before to remove the MBP tag. AVR-Pii was purified from the MBP solubility tag using HisTrap™ and MBPTrap™ (GE Healthcare) columns attached in tandem **(Figure S3b)**. Purified AVR-Pii was commonly present as a double band in the flow-through (FT) and wash-through (WT) from the columns **(Figure S3b)**.

The relevant fractions were concentrated and loaded onto a Superdex 75 16/60 gel filtration column for final purification and buffer exchange into 20 mM HEPES (pH 7.5), 150 mM NaCl and 5% (vol/vol) glycerol **(Figure S3c)**. Relevant fraction with purified AVR-Pii were concentrated as appropriate and used for structural and biophysical characterization.

The state of the protein was assessed by intact mass spectrometry, revealing a main peak with a molecular weight of 5677.68 Da, identical to that calculated for AVR-Pii **(Figure S4)**.

All protein concentrations were determined using a Direct Detect Infrared Spectrometer (Merck).

#### Crystallization, data collection and structure solution

For crystallization, OsExo70F2 (residues 84 to 689) in complex with AVR-Pii (residues 20 to 70) was concentrated to 6 mg/ml following gel filtration. Sitting drop vapor diffusion crystallization trials were set up in 96 well plates, using an Oryx nano robot (Douglas Instruments, United Kingdom), with drops comprised of 0.3 μl precipitant solution and 0.3 μl of the sample, and incubated at 20°C. After four to six days, protein crystals for the complex between OsExo70F2 and AVR-Pii appeared in the 0.3 M Ammonium iodide; 30% v/v PEG3350 condition of the PEG Suite screen (Qiagen). For data collection, all crystals were harvested using Litholoops (Molecular Dimensions) and flash-cooled in liquid nitrogen.

X-ray data were collected from a single crystal at the Diamond Light Source using beamline i03 (Oxford, UK) at 100 K and recorded on a Pilatus3 6M hybrid photon counting detector (Dectris). The data were processed using the xia2 pipeline (85) and CCP4 (86). To solve the structure of OsExo70F2/AVR-Pii complex, we used the *Arabidopsis* Exo70A1 (PDB ID: 4L5R) divided into three ensembles (Ensemble 1 residues 76 to 325; Ensemble 2 residues 326 to 474; Ensemble 3 residues 475 to 593) as a template for molecular replacement using PHASER (87). Once we obtained a solution, automated model building using BUCANNEER (88) was able to identify and build the AVR-Pii effector. The asymmetric unit of the crystal contains only a single copy of the OsExo70F2/AVR-Pii complex with a stoichiometry of 1:1. The final structure was obtained through iterative cycles of model building and refinement using COOT (Emsley et al., 2010), REFMAC5 (89), and ISOLDE (90) as implemented in the CCP4 suite (86) and ChimeraX (61). Structures were validated using the tools provided in COOT and MOLPROBITY (91).

#### Yeast-2-hybrid

OsExo70 proteins encoded in pGBKT7 plasmids were co-transformed with AVR-Pii variants or mutants in pGADT7 into chemically competent Y2HGold cells (Takara Bio, USA) using a Frozen-EZ Yeast Transformation Kit (Zymo research).

Single colonies grown on selection plates were inoculated in 5 ml of SD-Leu-Trp overnight at 30 °C. Saturated culture was then used to make serial dilutions of OD_600_ 1, 0.1, 0.01, and 0.001. 5 μl of each dilution was spotted on a SD-Leu-Trp plate as a growth control, and on a SD-Leu-Trp-Ade-His plate containing X-α-gal and supplemented with 0.2 or 1 μg/ml Aureobasidin A (Takara Bio, USA). Plates were imaged after incubation for 60–72 hr at 30 °C. Each experiment was repeated a minimum of three times, with similar results.

To confirm protein expression, total yeast extracts from transformed colonies were produced by boiling the cells for 10 min in LDS Runblue sample buffer. Samples were centrifugated and the supernatant was subjected to SDS-PAGE gels and western blot. The membranes were probed with anti-GAL4 DNA-BD (Sigma) for the OsExo70 proteins in pGBKT7 and with the anti-GAL4 activation domain (Sigma) antibodies for the AVR-Pii wild type and mutant effectors in pGADT7.

---

**Figure S1.**
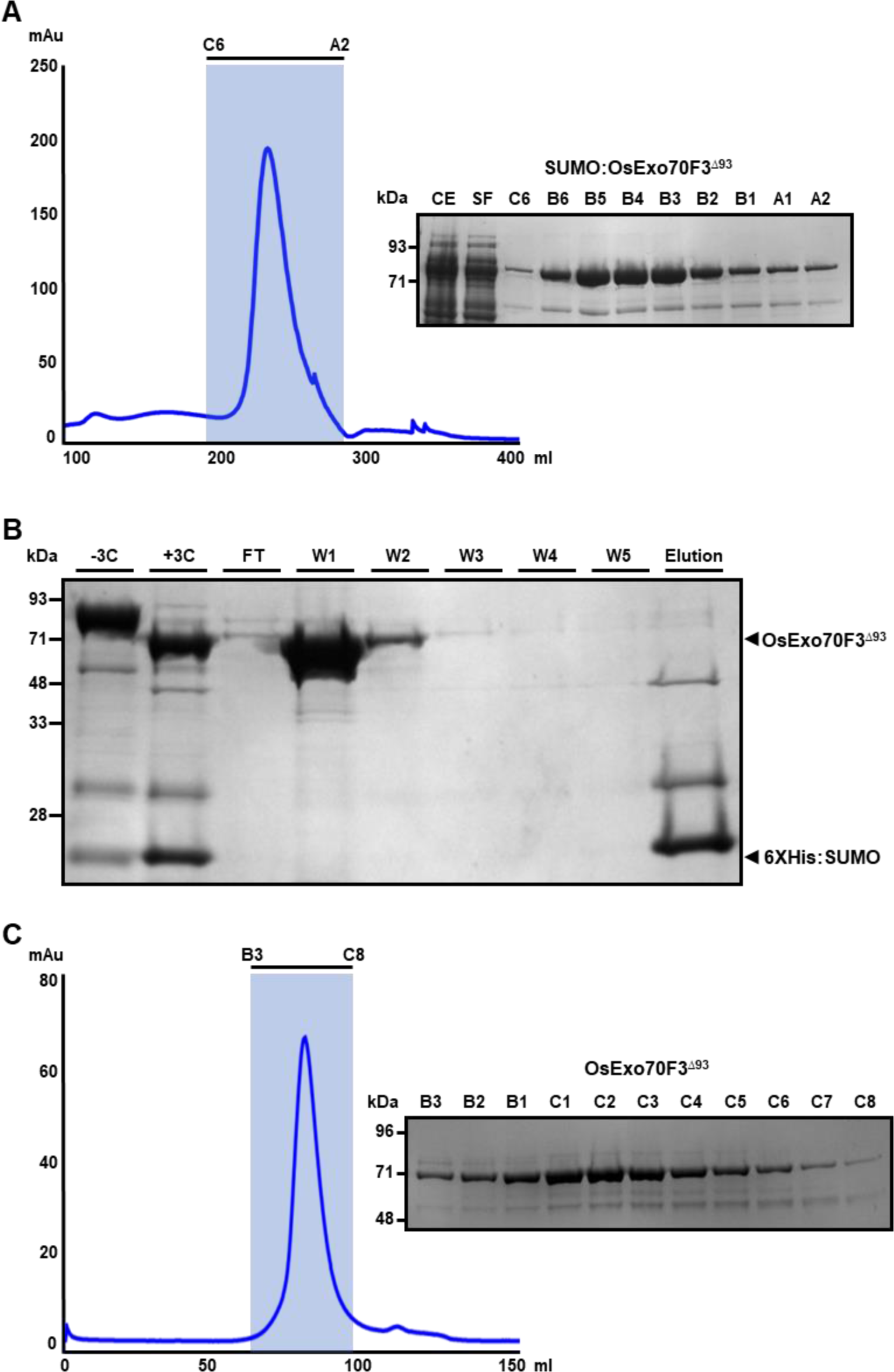
Expression and purification of rice OsExo70 proteins. **(A)** Elution trace of SUMO:OsExo70F3^Δ93^ after IMAC and gel filtration with selected fractions analysed by SDS-PAGE. **(B)** SDS-PAGE analysis of fractions collected in the OsExo70F3^Δ93^ HisTrap™ purification before and after cleaving the SUMO tag with 3C protease. OsExo70F3^Δ93^ was successfully purified in the first and second wash fractions. The 6xHis:SUMO tag is released upon treatment with elution buffer. **(C)** Elution trace of OsExo70F3^Δ93^ after gel filtration and SDS-PAGE analysis of relevant fractions.

**Figure S2.**
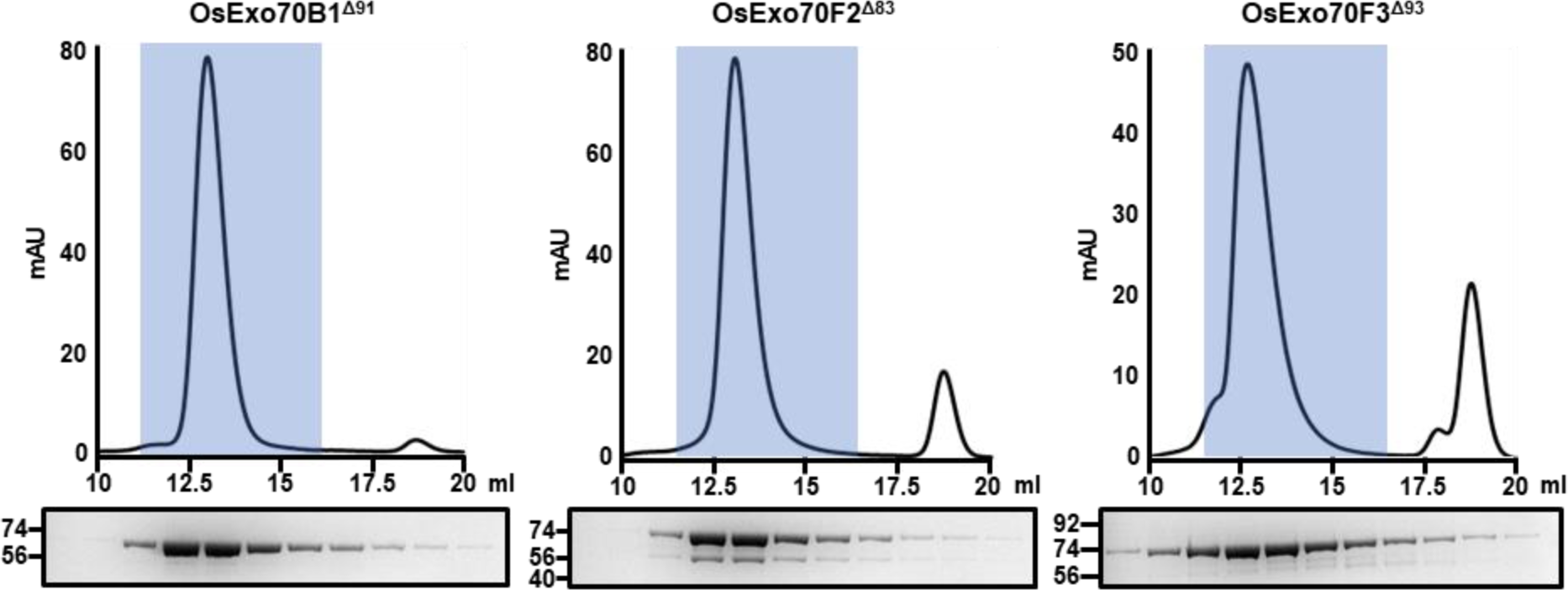
Purified plant Exo70 proteins elute as single peaks in analytical gel filtration analysis. Elution traces of purified OsExo70B1^Δ91^, OsExo70F2^Δ83^ and OsExo70F3^Δ93^ in analytical gel filtration. SDS-PAGE analysis of the fractions highlighted in ice blue are shown below each trace.

**Figure S3.**
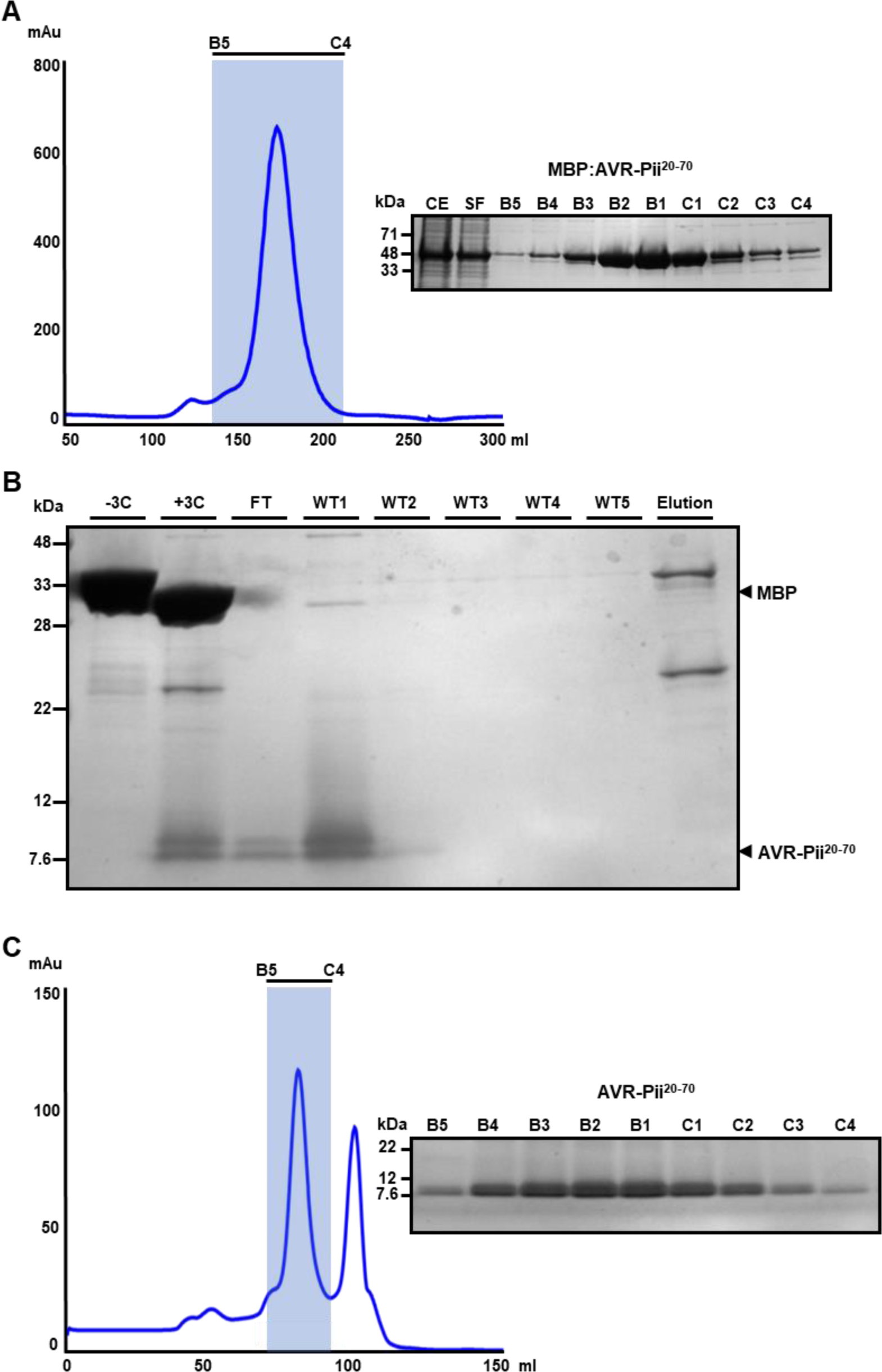
Expression and purification of AVR-Pii. **(A)** Elution trace of MBP:AVR- Pii^20-70^ after IMAC and gel filtration with selected fractions analysed by SDS-PAGE. **(B)** SDS-PAGE analysis of fractions collected in the AVR-Pii^20-70^ HisTrap™/ MBPTrap™ purification before and after cleaving the MBP tag with 3C protease. AVR-Pii^20-70^ was successfully purified in the flow-through and first wash fractions. **(C)** Elution trace of AVR-Pii^20-70^ after gel filtration and SDS-PAGE analysis of relevant fractions.

**Figure S4.**
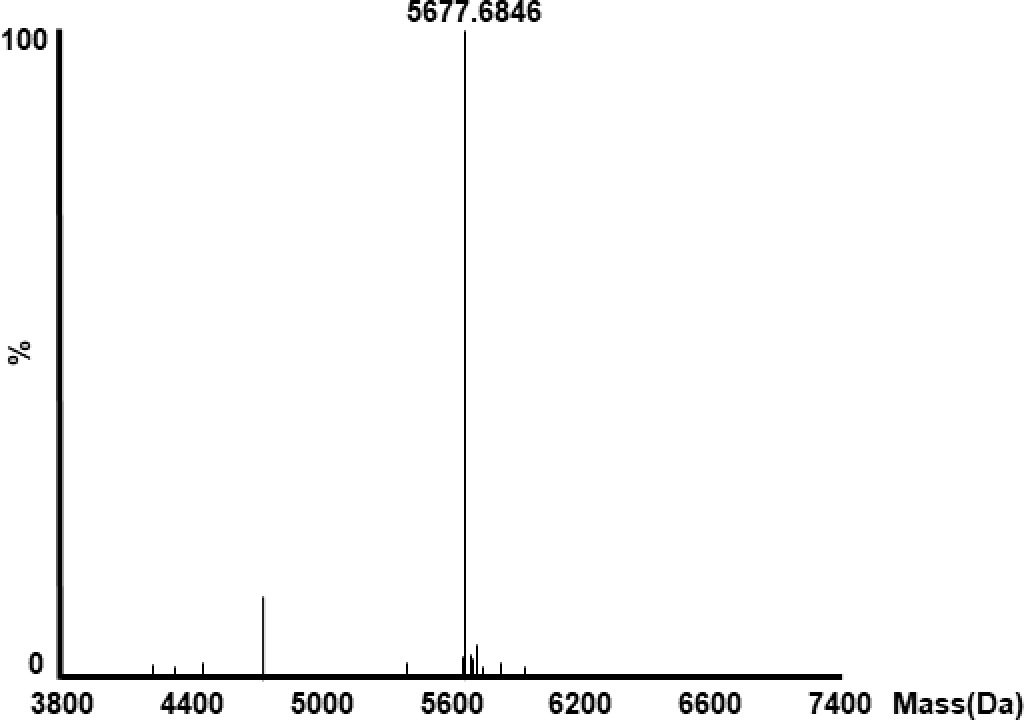
Mass spectrometry analysis of purified AVR-Pii. Intact mass analysis of purified AVR-Pii^20-70^ protein shows a main peak close to the predicted protein mass (5,675.0 Da).

**Figure S5.**
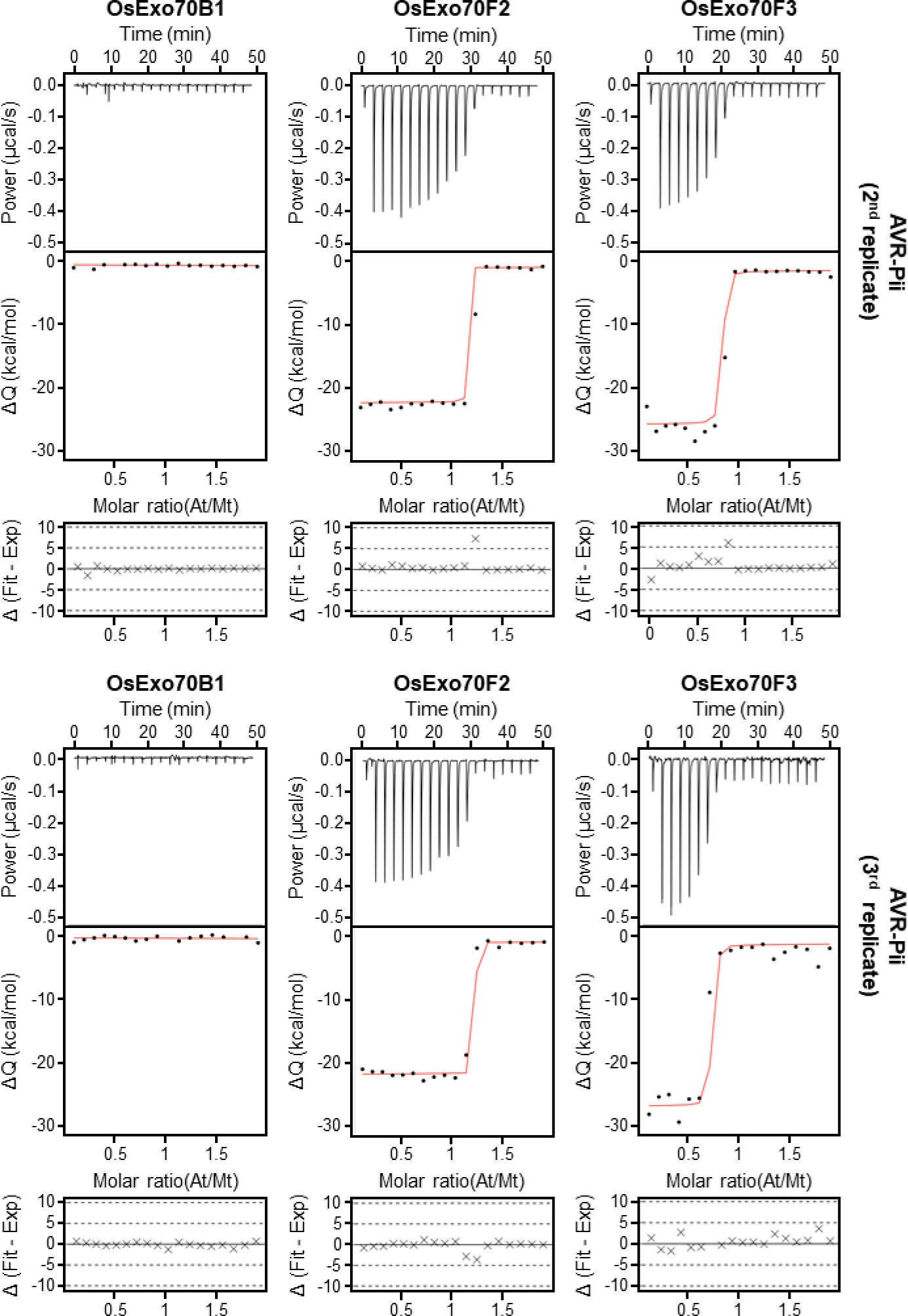
Experimental replicates of AVR-Pii binding to OsExo70s measured by ITC. Binding of AVR-Pii to rice OsExo70 proteins determined by isothermal titration calorimetry (ITC). Upper panels show heat differences upon injection of AVR-Pii into the cell containing the respective OsExo70 allele. Middle panels show integrated heats of injection (dots) and the best fit (solid line) using to a single site binding model calculated using AFFINImeter ITC analysis software (79). Bottom panels represent the difference between the fit to a single site binding model and the experimental data; the closer to zero indicates stronger agreement between the data and the fit. The thermodynamic parameters obtained in each experiment are presented in **Table S1**.

**Figure S6.**
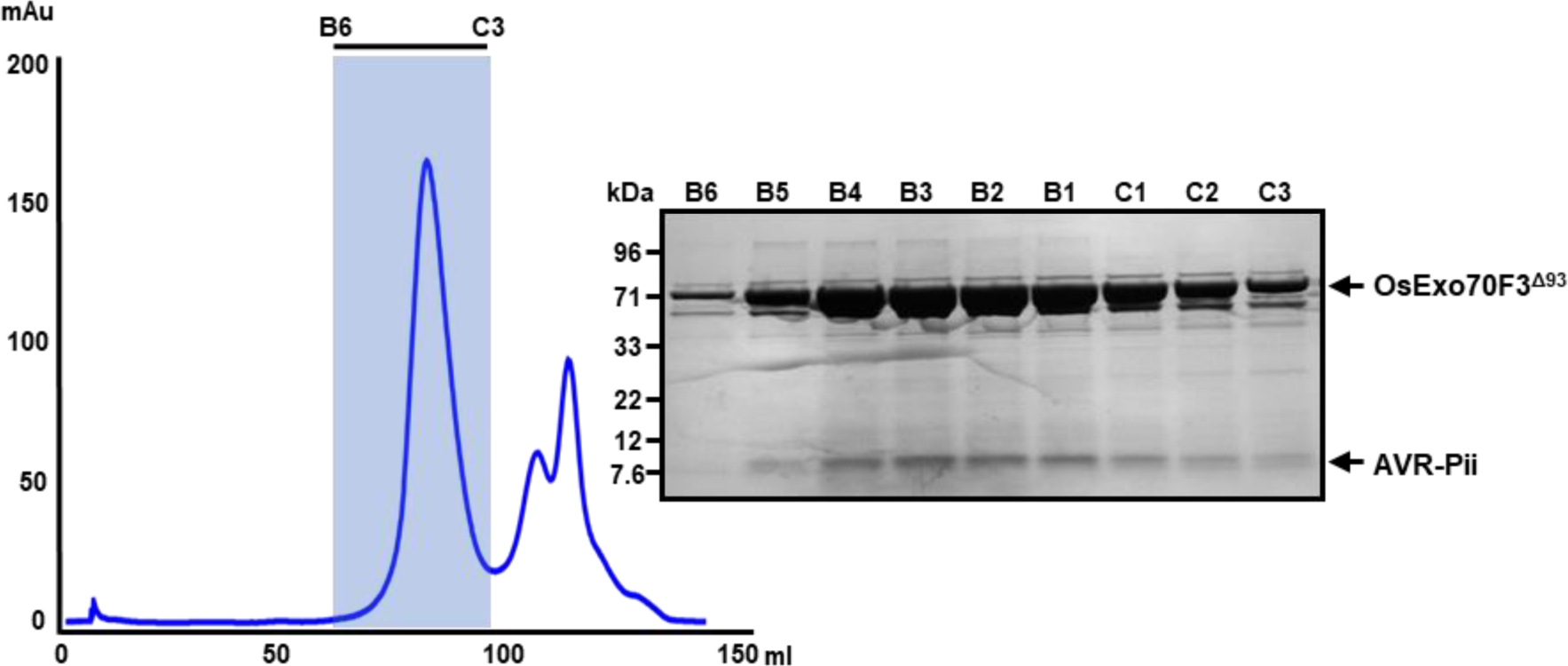
The OsExo70/AVR-Pii complex can be reconstituted in vitro. Elution trace of reconstitution of OsExo70/AVR-Pii complex after gel filtration using OsExo70F3^Δ93^ and AVR-Pii as example. Selected fractions were collected and analysed by SDS-PAGE showing the presence of both proteins.

**Figure S7.**
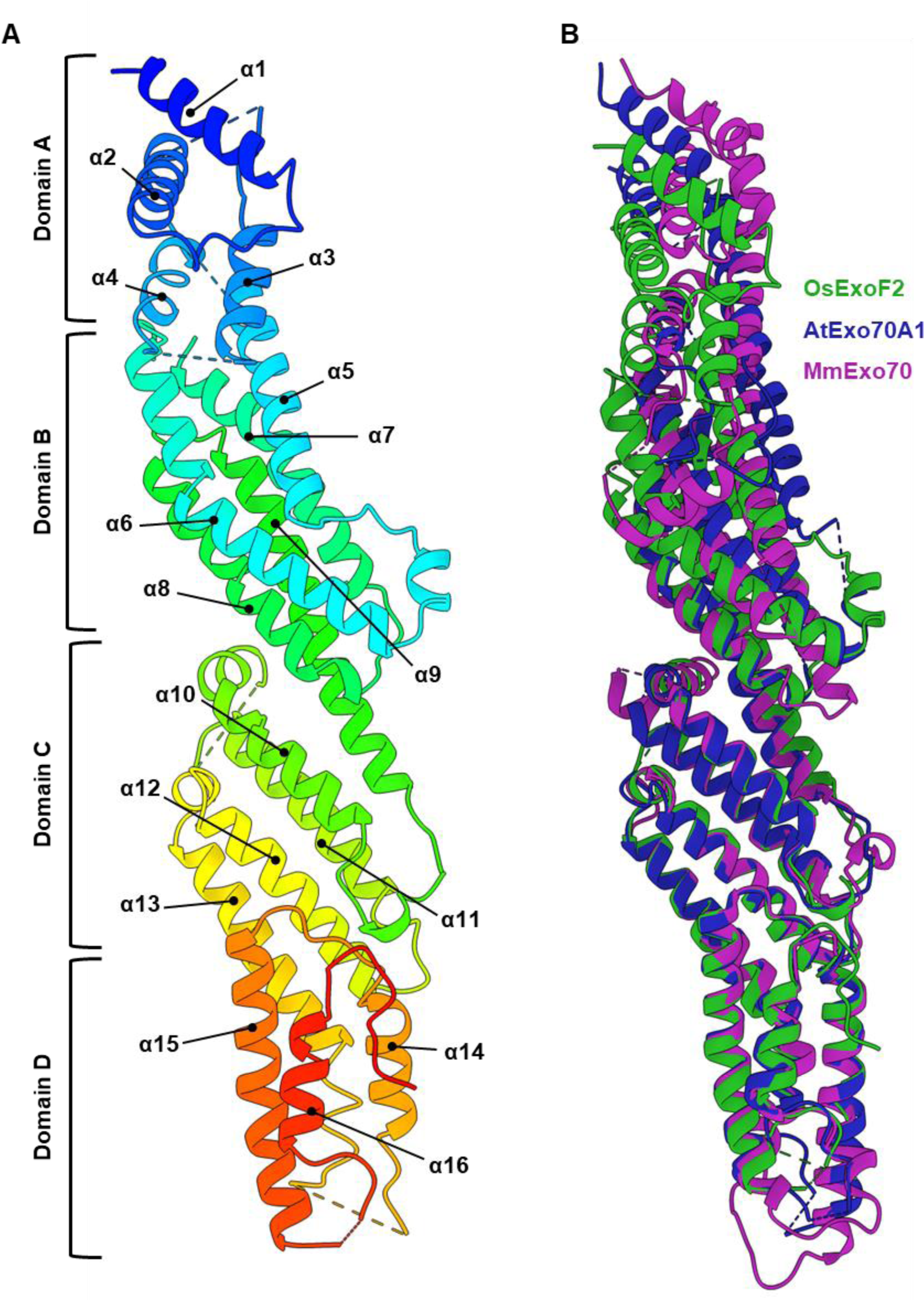
OsExo70F2 adopts a conserved Exo70 fold. **(A)** Schematic representation of OsExo70F2 domains with α-helices as cartoon ribbons. **(B)** Superposition of the overall structures of rice OsExo70F2 (green), Arabidopsis AtExo70A1 (PDB ID: 4L5R) (57) (dark blue) and mouse Exo70 (PDB ID: 2PFT) (56) (purple). Each Exo70 is coloured as labelled and is shown in cartoon representation.

**Figure S8.**
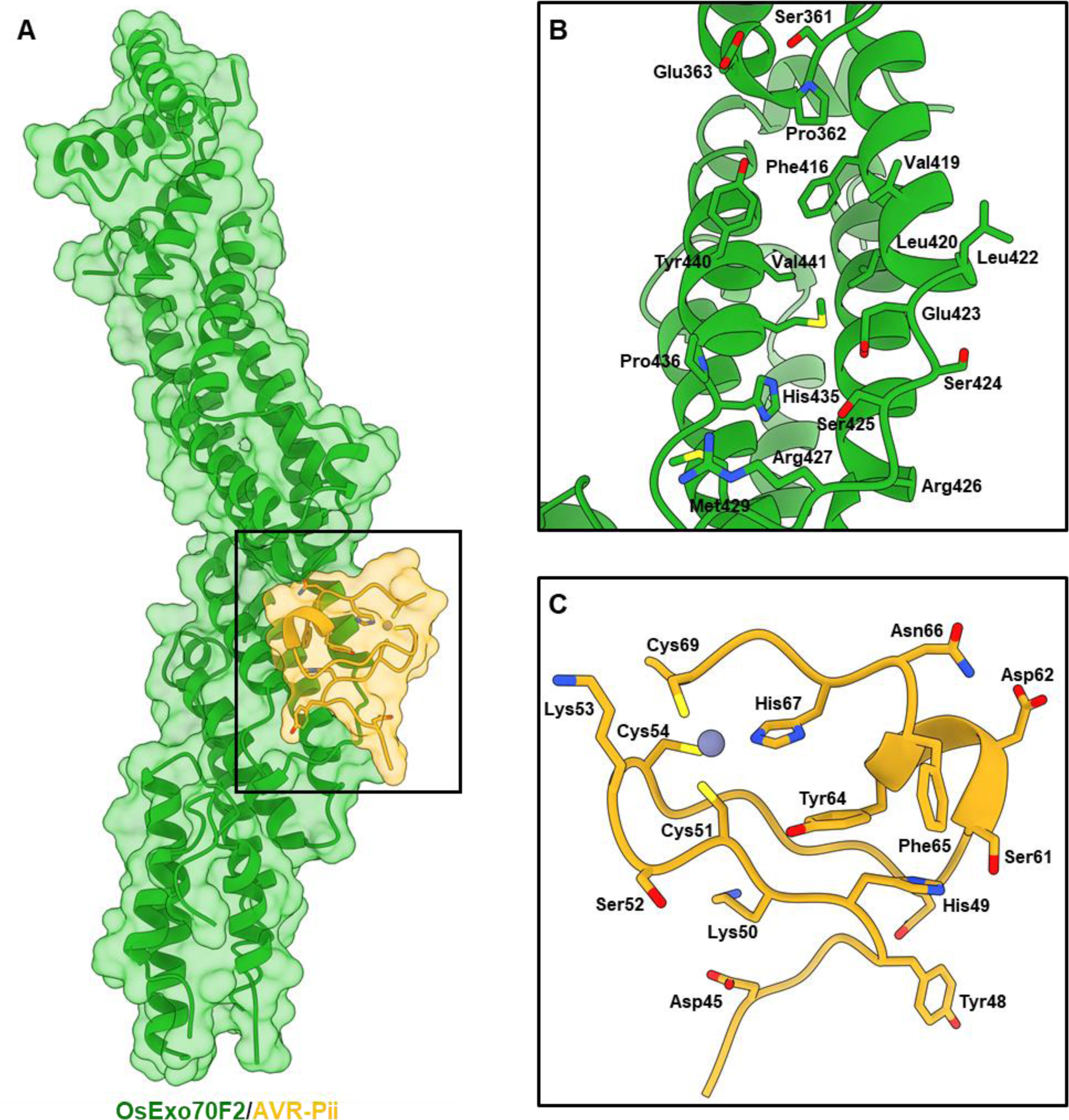
Detailed view of residues at the OsExo70F2/AVR-Pii interface. **(A)** Schematic representation of OsExo70F2/AVR-Pii complex represented as cartoon ribbons with the molecular surface also shown and coloured as labelled. The interaction interface is delimited by the black square. **(B)** Close-up view of residues comprising the OsExo70F2 interaction interface with AVR-Pii represented as cartoon ribbons. Residues forming the interaction interface are labelled with their side-chains displayed as cylinders. **(C)** Close-up view of residues comprising the AVR-Pii interaction interface with OsExo70F2 represented as cartoon ribbons. Residues forming the interaction interface are labelled with their side-chains displayed as cylinders.

**Figure S9.**
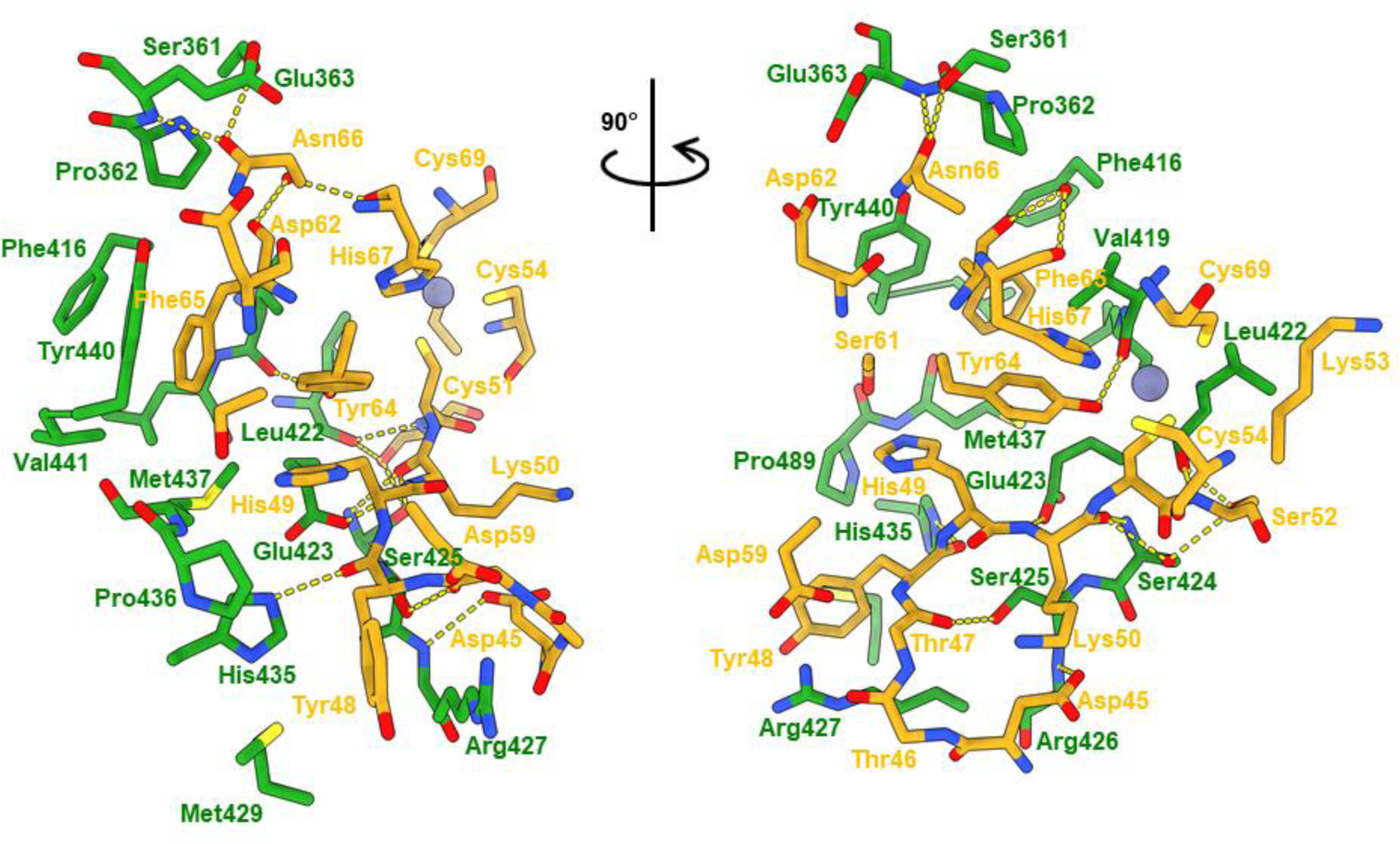
Interfacing residues within the OsExo70F2/AVR-Pii complex. Residues at OsExo70F2/AVR-Pii interaction interface displayed as cylinders. Residues from OsExo70F2 and AVR-Pii are coloured green or yellow, respectively. Hydrogen bonds/salt bridges are shown as yellow dashed lines.

**Figure S10.**
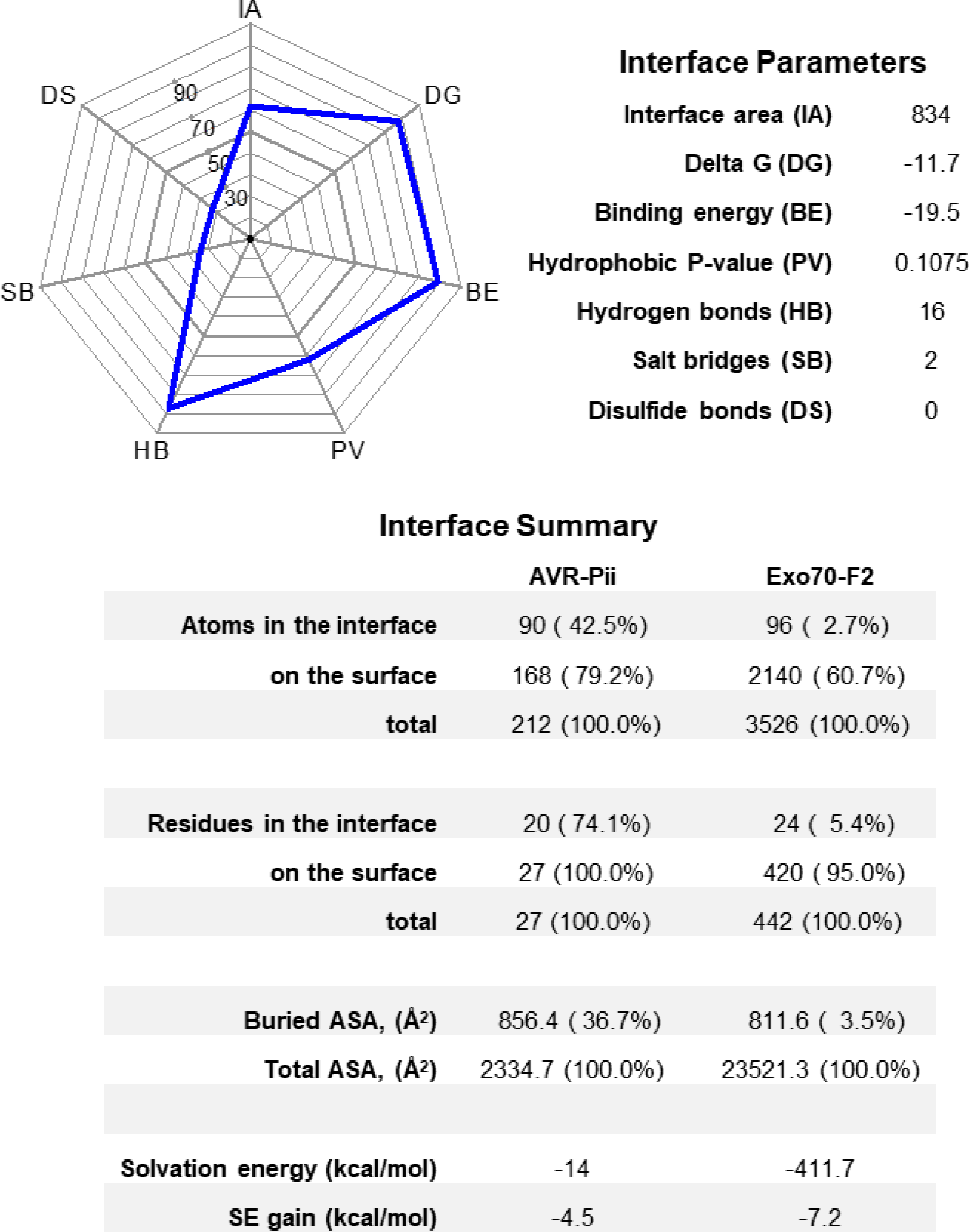
Analysis of the binding interface between OsExo70F2 and AVR-Pii using qtPISA. Interface analysis was performed using qtPISA (63). The key interface parameters in the analysis are represented as an interaction radar and the values are listed in the adjacent table.

**Figure S11.**
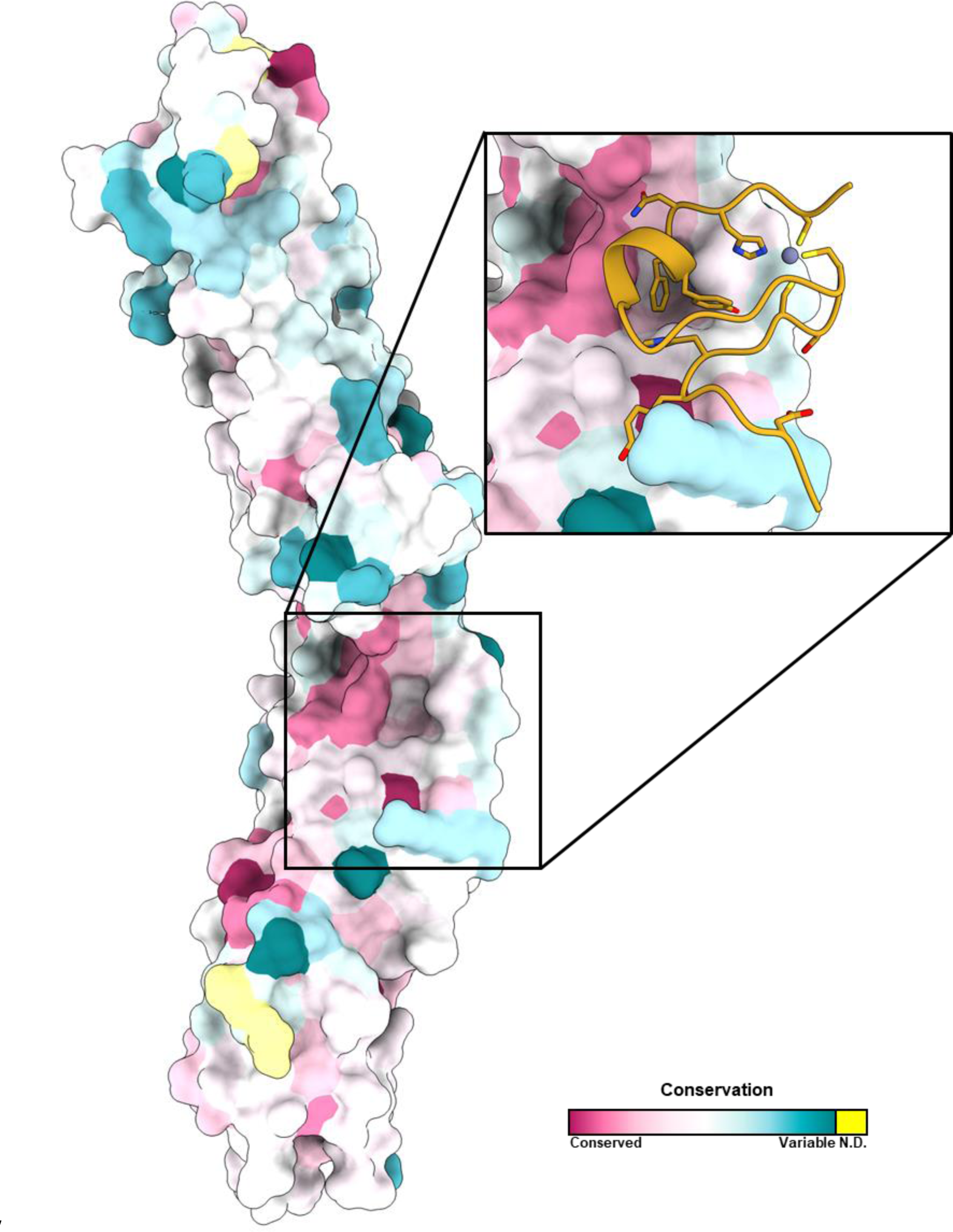
The amphipathic AVR-Pii binding interface is conserved in rice Exo70s. Conservation profile of rice Exo70 residues with a close-up view at the AVR-Pii interface as calculated by ConSurf (64). Exo70 is represented with solid surface coloured according to the conservation of their residues ranging from purple (highly conserved) through white (moderately conserved) to cyan (highly variable). Surface areas highlighted in yellow correspond to residues for which a meaningful conservation level could not be derived from the set of homologues used for the analysis. A close-up view of the effector interface is also shown with AVR-Pii represented in ribbons and coloured in yellow with the side chains of important residues displayed as cylinders.

**Figure S12.**
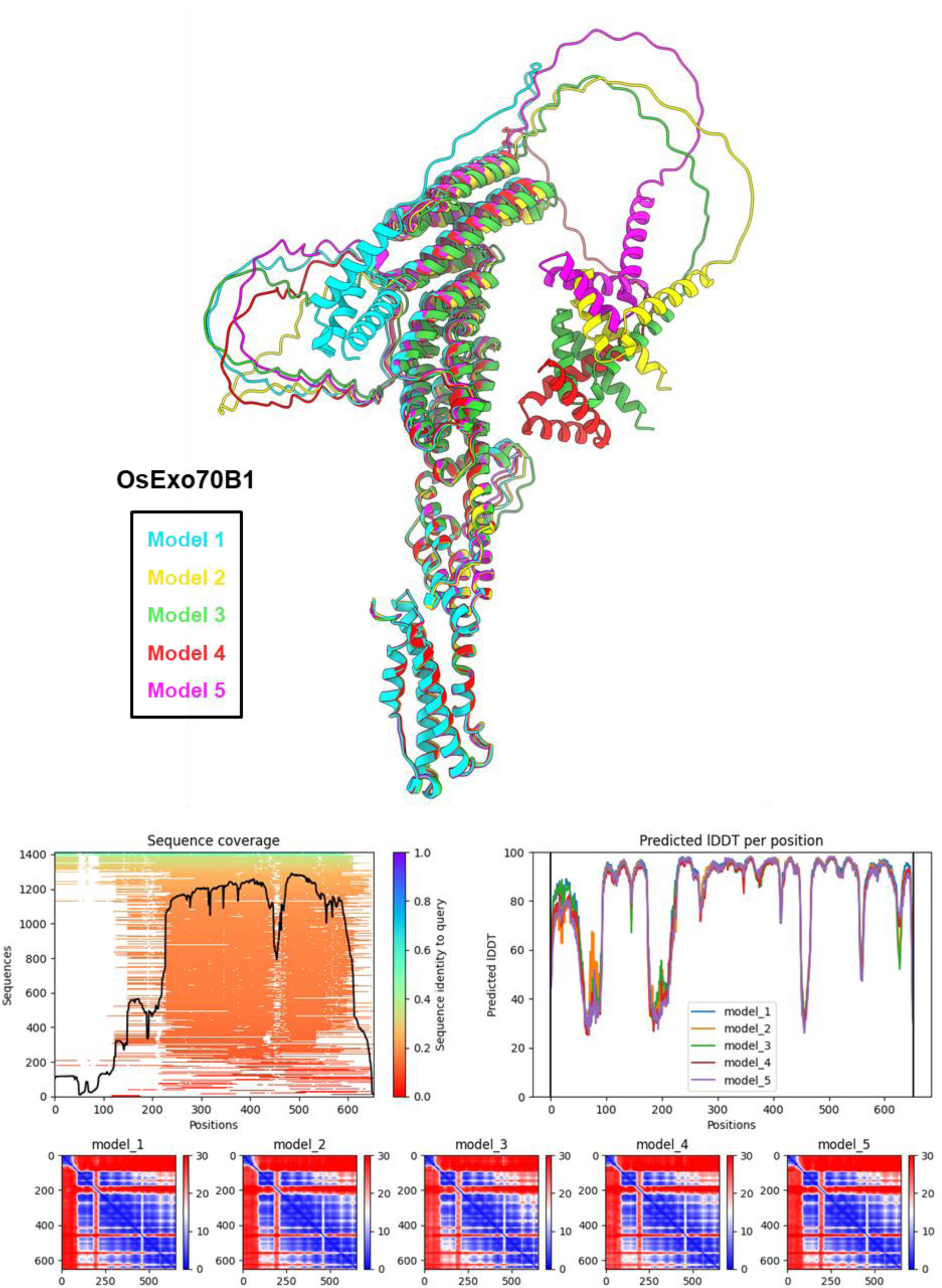
Homology modelling of OsExo70B1 calculated with AlphaFold2. Structure of OsExo70B1 generated by homology modelling using AlphaFold2 (65) as implemented by ColabFold (66).

**Figure S13.**
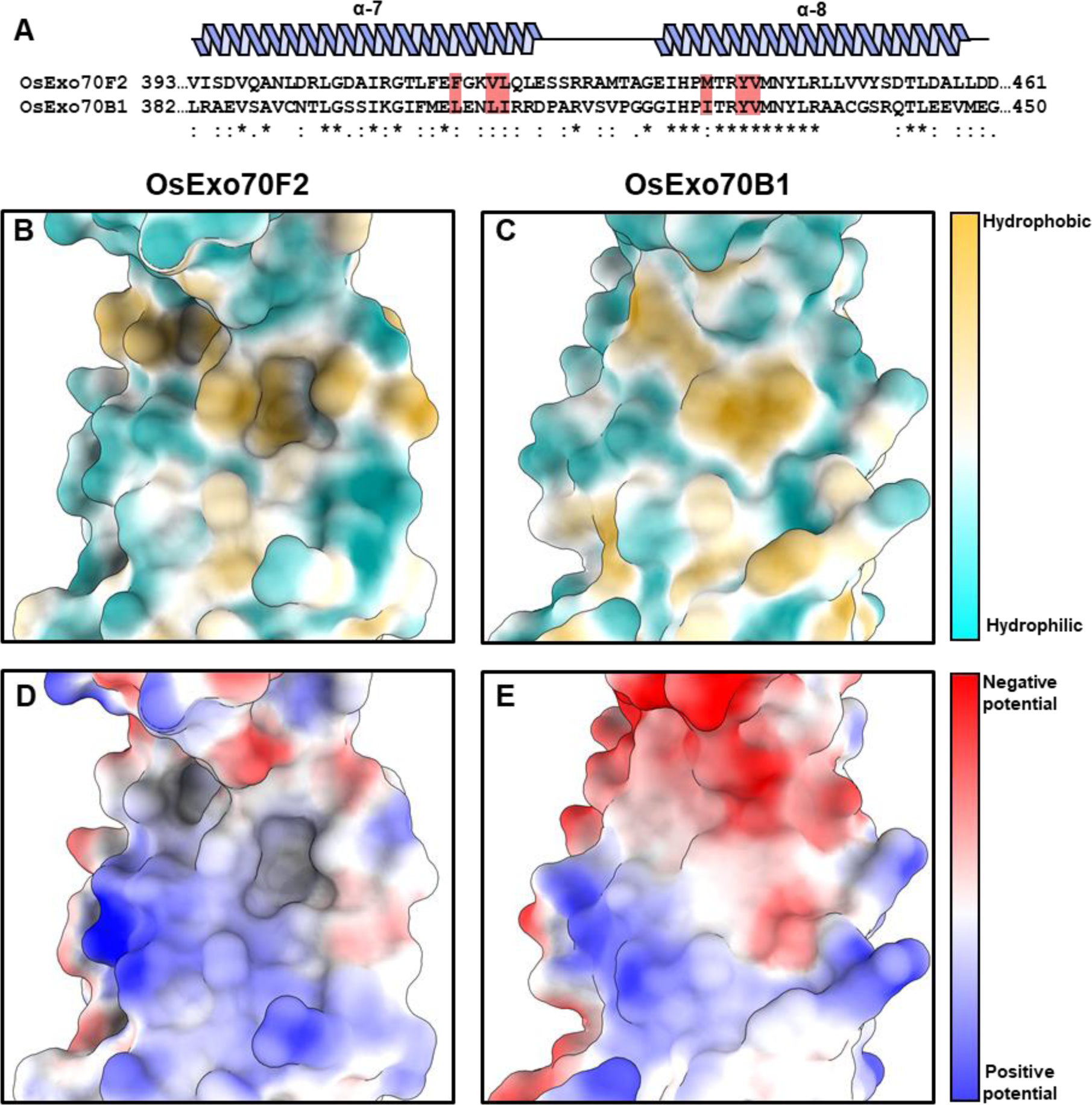
Comparison of the effector binding interface between OsExo70F2 and OsExo70B1. **(A)** Sequence alignment of residues located at the OsExo70F2 and OsExo70B1 α-helices 7 and 8 generated with Clustal Omega (92). Secondary structure features of Exo70 fold are shown above, and important residues for the formation of the AVR-Pii binding pocket are highlighted in red. Comparison of **(B)** OsExo70F2 and **(C)** OsExo70B1 surface hydrophobicity at the interaction interface with AVR-Pii, residues are coloured depending on their hydrophobicity from light blue (low) to yellow (high). Comparison of **(D)** OsExo70F2 and **(E)** OsExo70B1 surface electrostatic potentials at the interaction interface with AVR-Pii, residues are coloured depending on their electrostatic potential from dark blue (positive) to red (negative). The OsExo70B1 structure used for comparison was generated using AlphaFold2 **(Figure S12)**.

**Figure S14.**
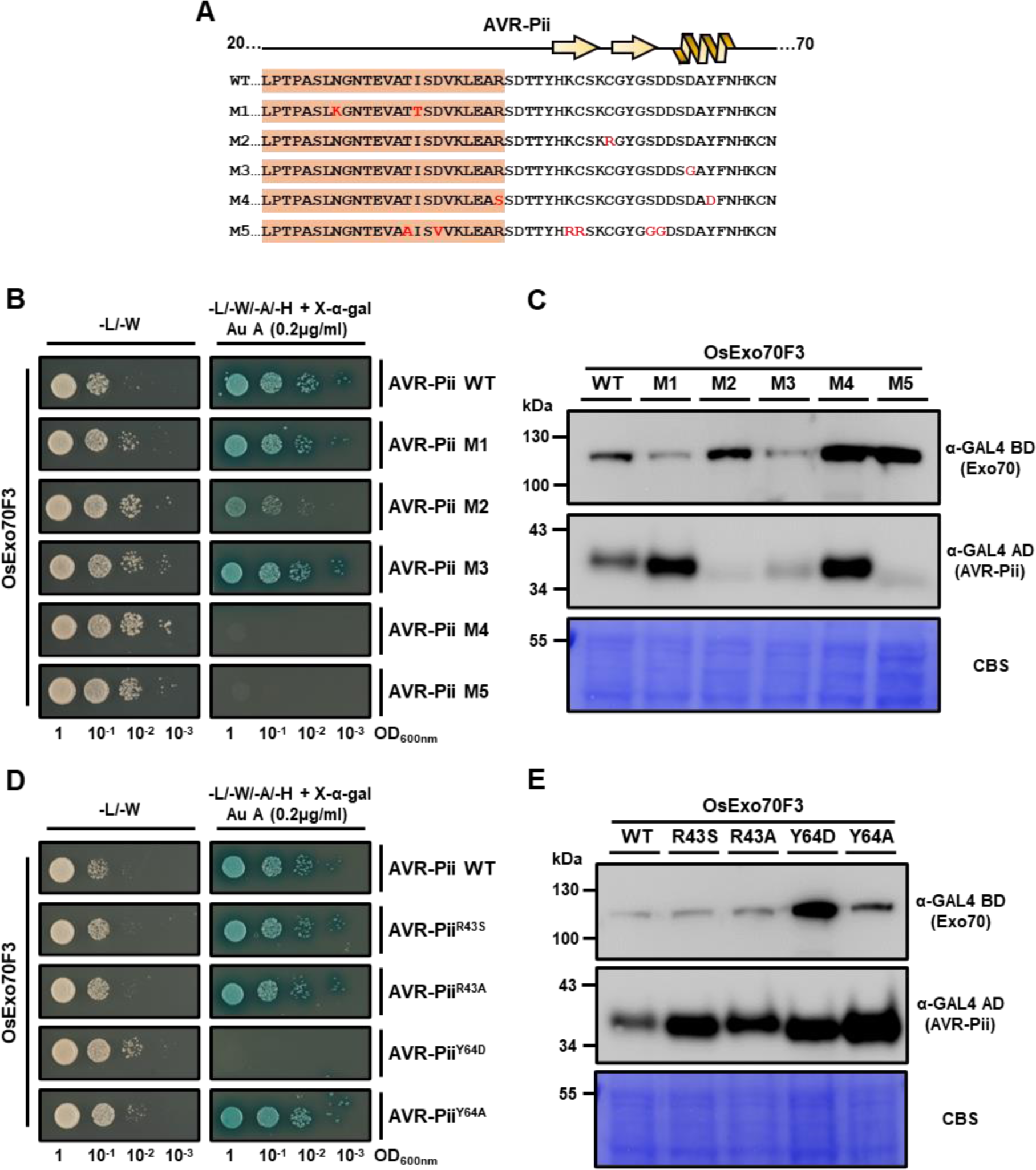
Random mutagenesis identified AVR-Pii residues that alter binding to OsExo70F3. **(A)** Amino acid sequence of the AVR-Pii mutants obtained by random mutagenesis. Secondary structure features of the AVR-Pii fold are shown above, and the residues not observed in the crystal structure are highlighted in orange. **(B)** Yeast-Two-Hybrid assay of AVR-Pii mutants obtained by random mutagenesis with rice Exo70F3. The control plate for yeast growth is on the left, with quadruple dropout media supplemented with X-α-gal and Aureobasidine A (Au A) on the right. Growth and development of blue colouration in the selection plate are both indicative of protein:protein interactions. Wild-type AVR-Pii is included as positive control. OsExo70F3 was fused to the GAL4 DNA binding domain, and AVR-Pii mutants to the GAL4 activator domain. Each experiment was repeated a minimum of three times, with similar results. **(C)** Accumulation of AVR-Pii mutants in yeast-two-hybrid assays analysed by Western blot. Yeast lysate was probed for the expression of rice Exo70F3 and AVR-Pii mutants obtained by random mutagenesis using anti-GAL4 binding domain (BD) and anti-GAL4 DNA activation domain (AD) antibodies, respectively. Total protein extracts were coloured with Coomassie Blue Stain (CBS). **(D)** Deconvolution of residues involved in AVR-Pii binding by Yeast-Two-Hybrid assay with rice Exo70F3. The control plate for yeast growth is on the left, with quadruple dropout media supplemented with X-α-gal and Aureobasidine A (Au A) on the right. Growth and development of blue colouration in the selection plate are both indicative of protein:protein interactions. Wild-type AVR-Pii is included as positive control. OsExo70F3 was fused to the GAL4 DNA binding domain, and AVR-Pii point mutants to the GAL4 activator domain. Each experiment was repeated a minimum of three times, with similar results. **(E)** Accumulation of AVR-Pii point mutants in yeast-two- hybrid assays analysed by Western blot. Yeast lysate was probed for the expression of rice Exo70F3 and AVR-Pii point mutants using anti-GAL4 binding domain (BD) and anti-GAL4 DNA activation domain (AD) antibodies, respectively. Total protein extracts were coloured with Coomassie Blue Stain (CBS).

**Figure S15.**
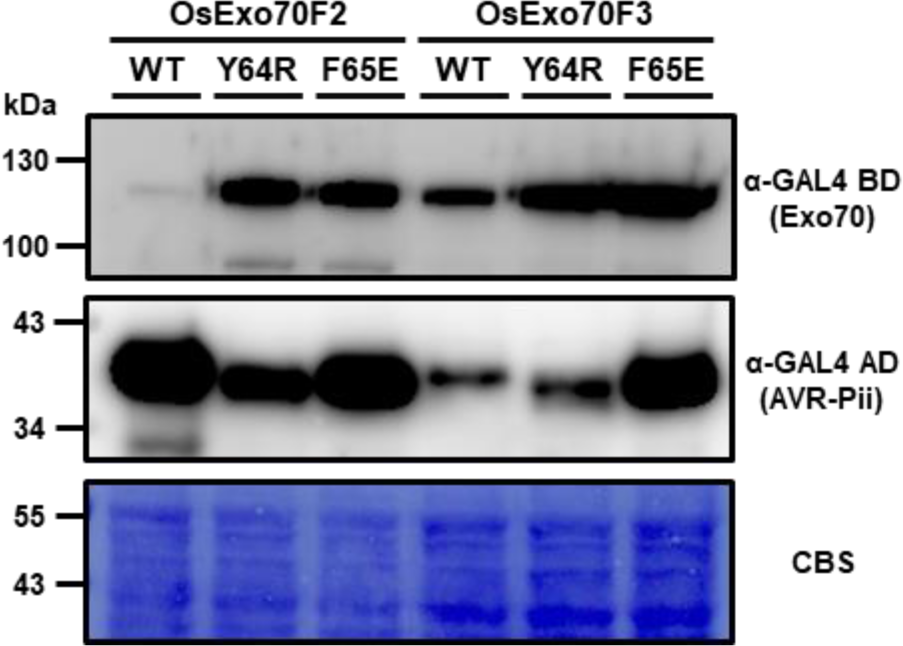
Accumulation of proteins in yeast-two-hybrid assays analysed by Western blot. Yeast lysate was probed for the expression of rice Exo70 proteins and AVR-Pii wild-type, AVR-Pii Tyr64Arg and AVR-Pii using anti-GAL4 binding domain (BD) and anti-GAL4 DNA activation domain (AD) antibodies, respectively. Total protein extracts were coloured with Coomassie Blue Stain (CBS).

**Figure S16.**
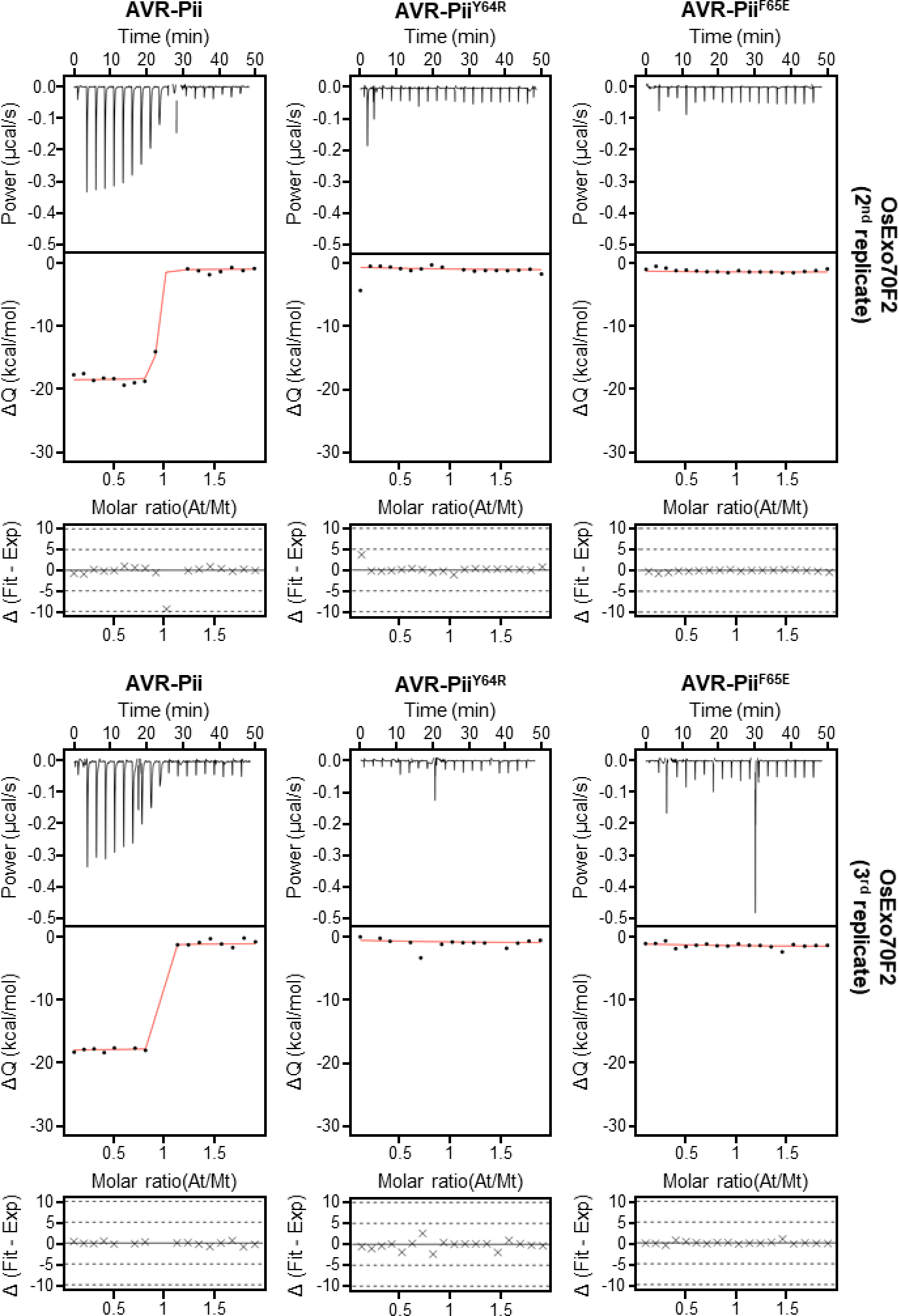
Experimental replicates for the binding of AVR-Pii mutants to OsExo70F2 measured by ITC. Upper panels show heat differences upon injection of AVR-Pii mutants into the cell containing OsExo70F2. Middle panels show integrated heats of injection (dots) and the best fit (solid line) using to a single site binding model calculated using AFFINImeter ITC analysis software (79). Bottom panels represent the difference between the fit to a single site binding model and the experimental data; the closer to zero indicates stronger agreement between the data and the fit. The thermodynamic parameters obtained in each experiment are presented in **Table S3**.

**Figure S17.**
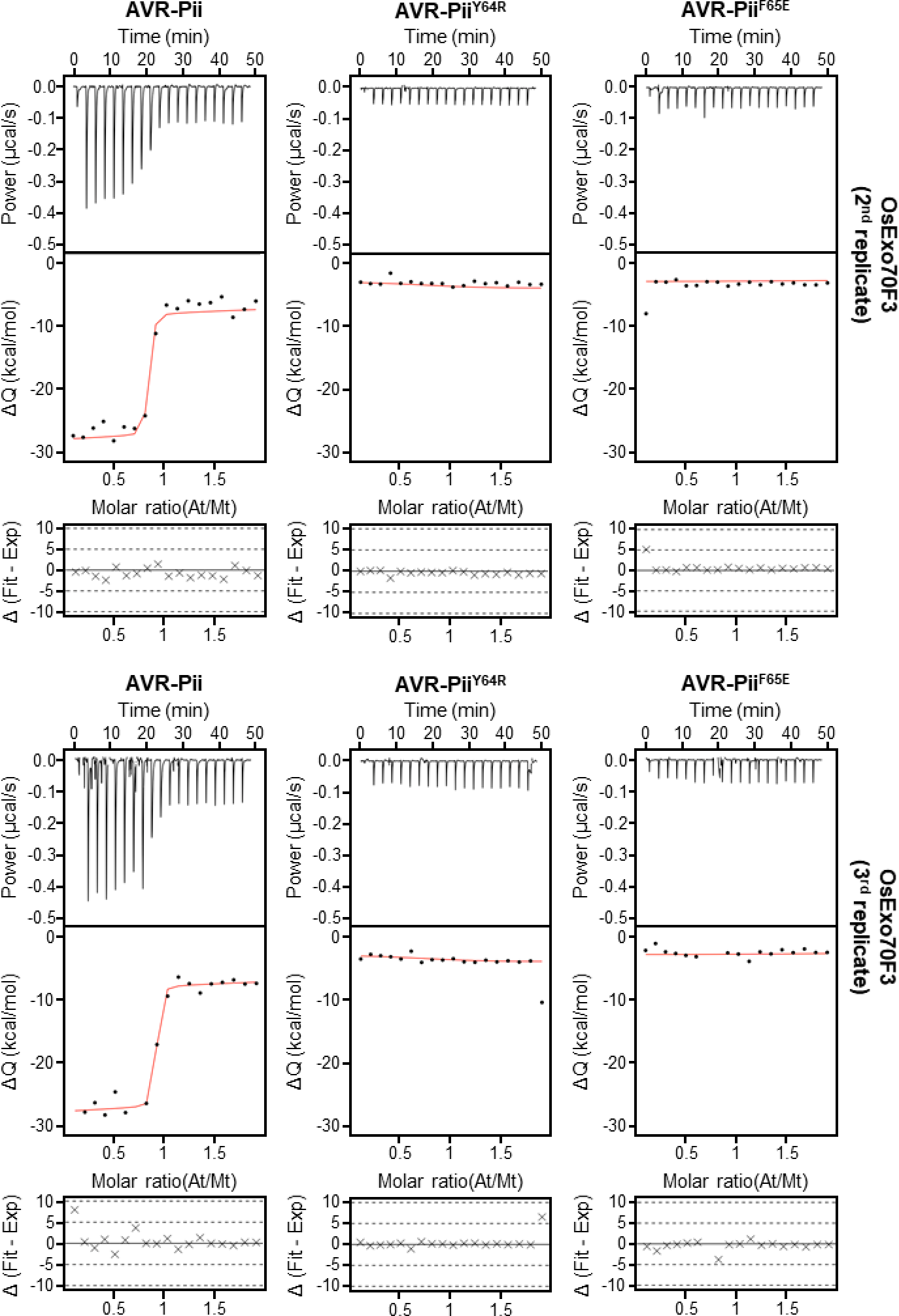
Experimental replicates for the binding of AVR-Pii mutants to OsExo70F3 measured by ITC. Upper panels show heat differences upon injection of AVR-Pii mutants into the cell containing OsExo70F3. Middle panels show integrated heats of injection (dots) and the best fit (solid line) using to a single site binding model calculated using AFFINImeter ITC analysis software (79). Bottom panels represent the difference between the fit to a single site binding model and the experimental data; the closer to zero indicates stronger agreement between the data and the fit. The thermodynamic parameters obtained in each experiment are presented in **Table S3**.

**Figure S18.**
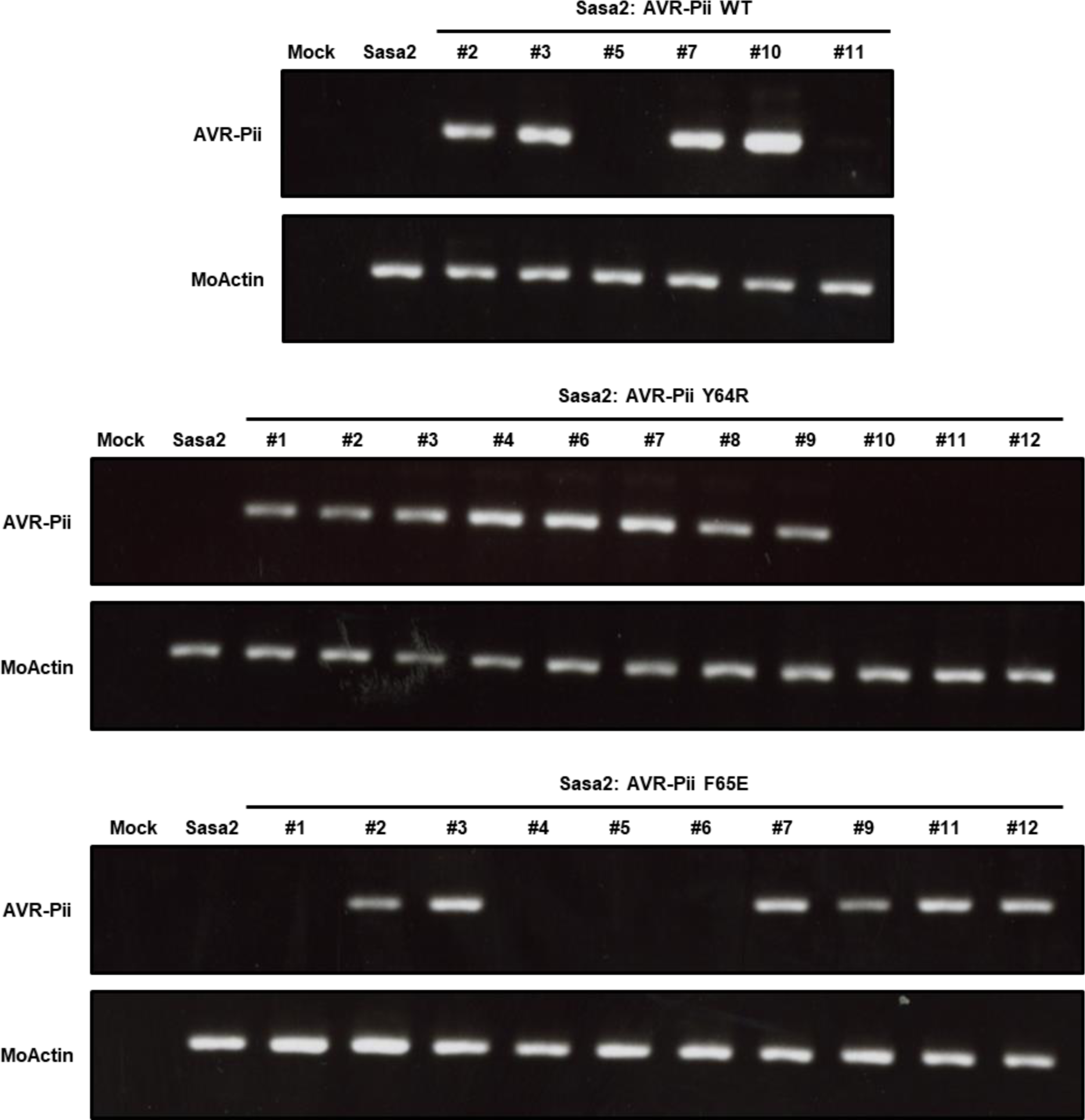
RT-PCR analysis of effector expression in Sasa2 isolates transformed with AVR-Pii and mutants. The presence of a band depicts that the gene is expressed in rice leaves during infection by the fungal isolates. The top panel corresponds to the expression of AVR-Pii WT or mutants. The expression of MoActin was tested as a positive control and is shown in the bottom panel.

**Figure S19.**
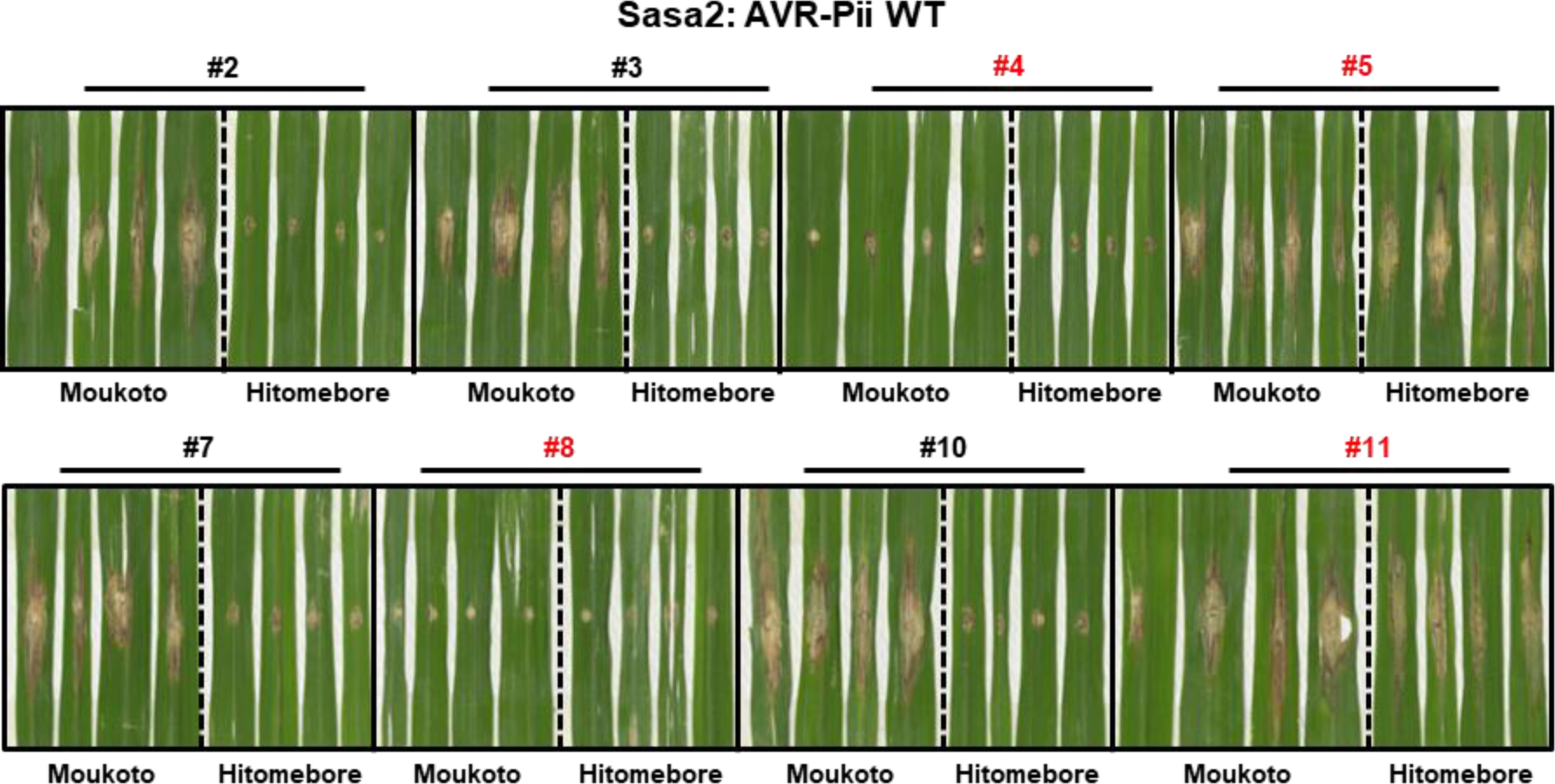
Replicates of the disease resistance assays of Sasa2 harbouring AVR-Pii WT. Rice leaf blade spot inoculation of transgenic *M. oryzae* Sasa2 into rice cultivars Moukoto (Pii-) and Hitomebore (Pii+). Eight independent Sasa2 transformants harbouring wild-type AVR-Pii were spotted in both cultivars. Line numbers coloured in red indicate the transformants removed from further quantification because they did not express AVR-Pii effectors as tested by RT-PCR **(Figure S18)** or were not infective.

**Figure S20.**
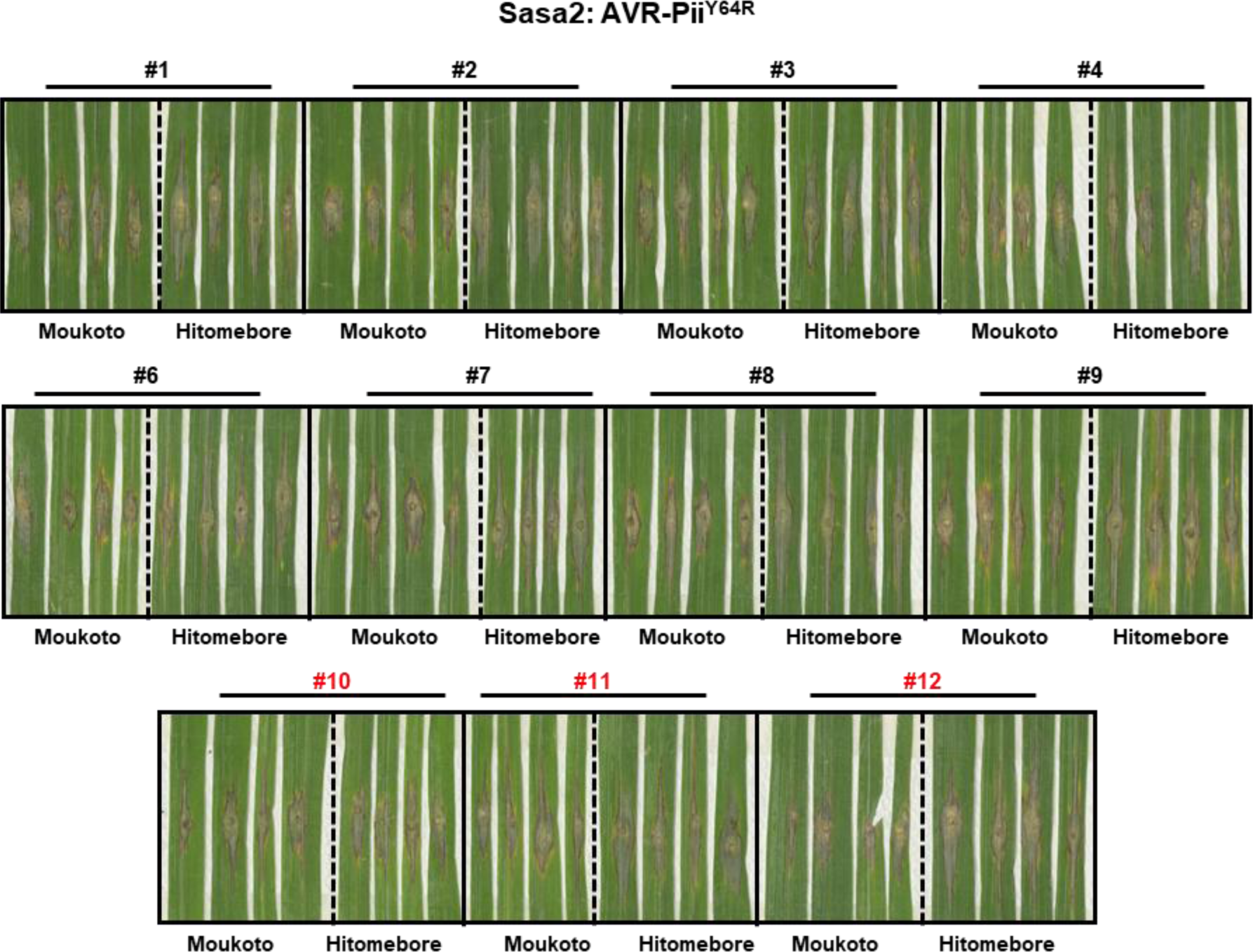
Replicates of the disease resistance assays of Sasa2 harbouring AVR-Pii Tyr64Arg. Rice leaf blade spot inoculation of transgenic *M. oryzae* Sasa2 into rice cultivars Moukoto (Pii-) and Hitomebore (Pii+). Eleven independent Sasa2 transformants harbouring mutant AVR-Pii Tyr64Arg were spotted in both cultivars. Line numbers coloured in red indicate the transformants removed from further quantification because they did not express AVR-Pii effectors as tested by RT-PCR **(Figure S18)** or were not infective.

**Figure S21.**
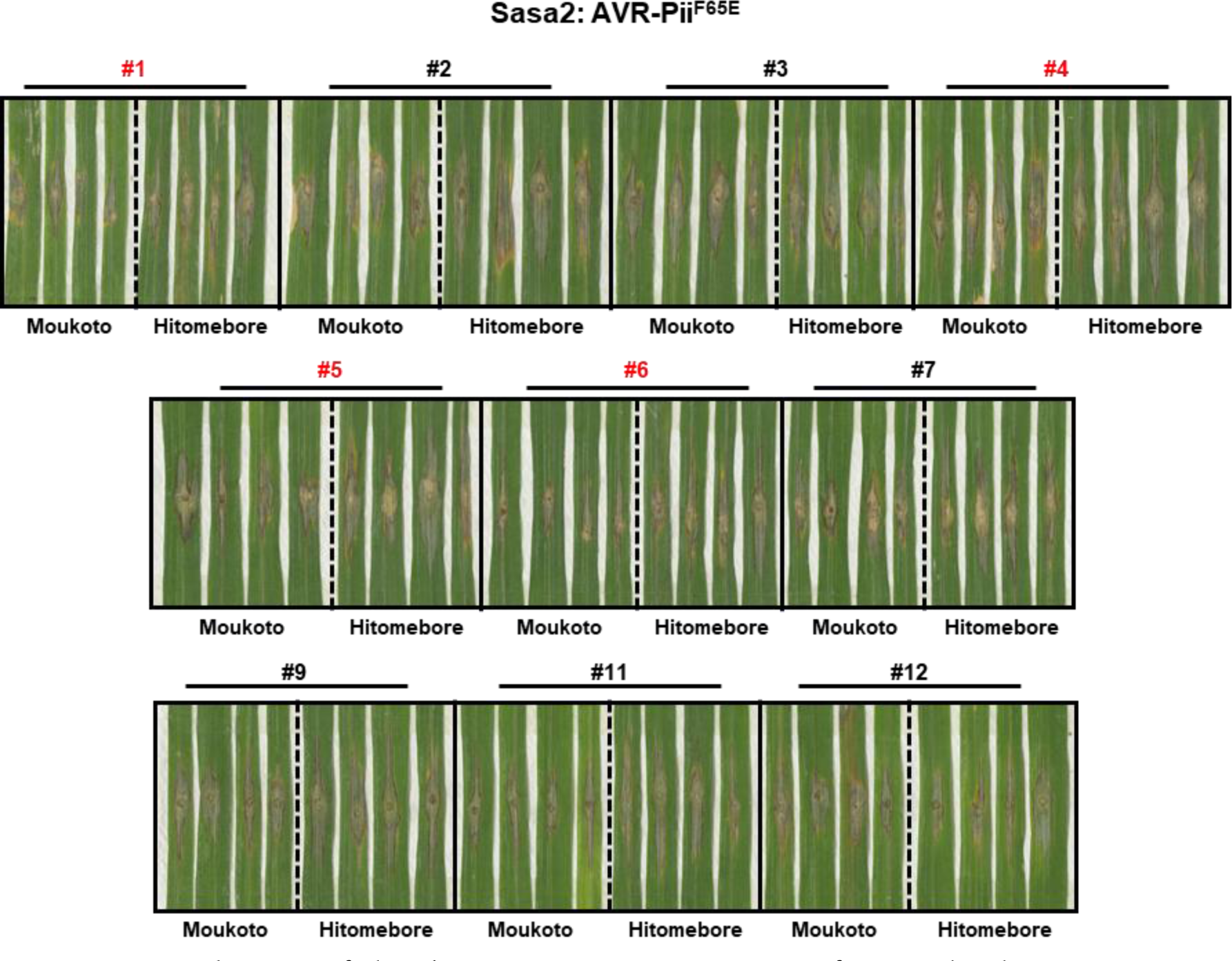
Replicates of the disease resistance assays of Sasa2 harbouring AVR-Pii Phe65Glu. Rice leaf blade spot inoculation of transgenic *M. oryzae* Sasa2 into rice cultivars Moukoto (Pii-) and Hitomebore (Pii+). Ten independent Sasa2 transformants harbouring mutant AVR-Pii Phe65Glu were spotted in both cultivars. Line numbers coloured in red indicate the transformants removed from further quantification because they did not express AVR-Pii effectors as tested by PCR **(Figure S18)** or were not infective.

**Figure S22.**
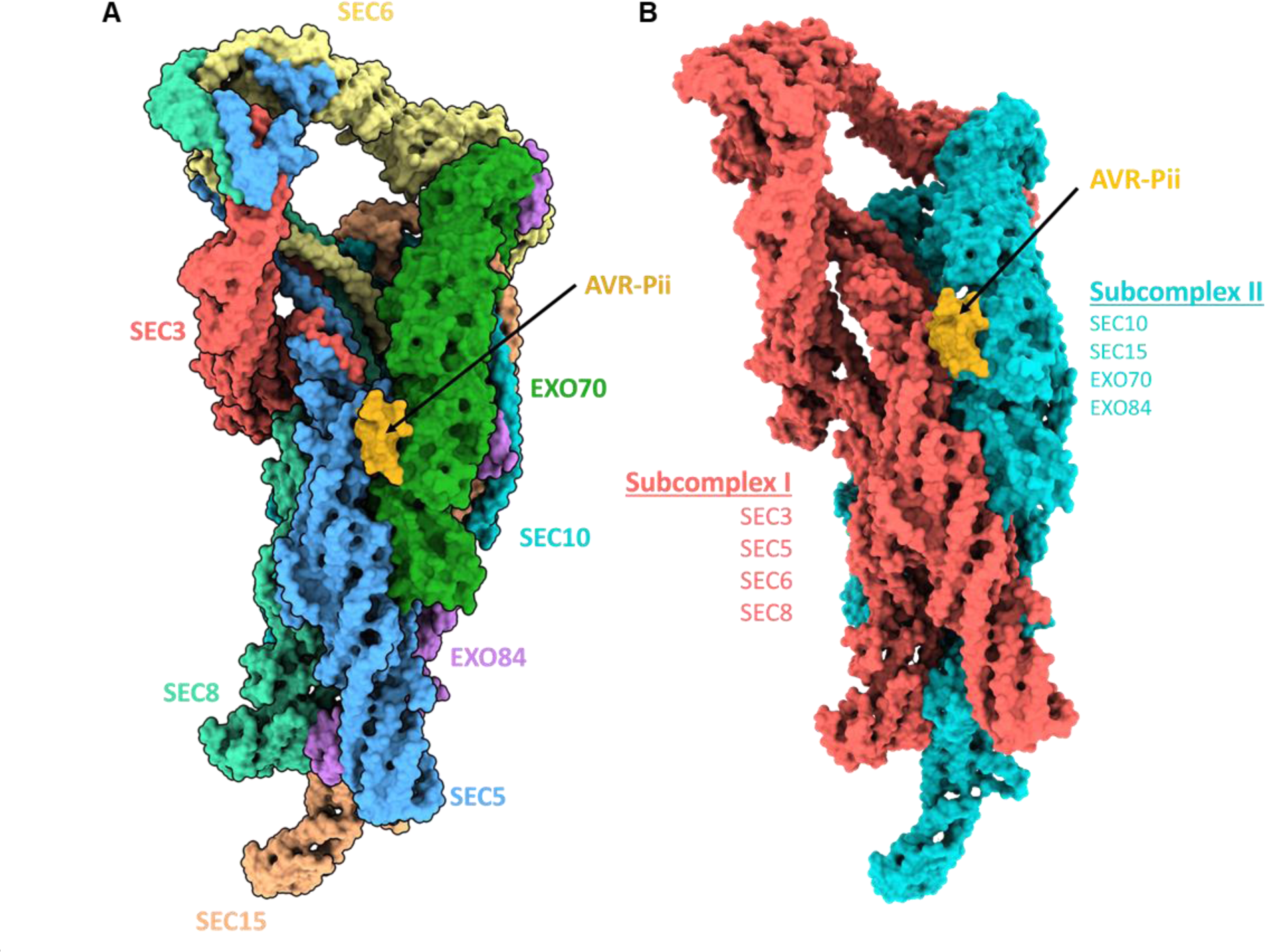
Representation of AVR-Pii interaction in the context of the exocyst complex. Structural alignment of AVR-Pii/OsExo70F2 structure in the Cryo-EM model of the yeast exocyst complex (PDB ID: 5YFP). The effector is coloured in gold and pointed by an arrow while the exocyst complex subunits are **(A)** individually coloured or **(B)** coloured according to the organization in subcomplexes.

**Table S1.** Experimental details and thermodynamic parameters obtained from ITC experiments presented in Figure 1.

| Cell | Conc. [$\mu\text{M}$] | Syringe | Conc. [$\mu\text{M}$] | T ($^{\circ}\text{C}$) | $\Delta\text{H}$ (kcal mol$^{-1}$) | $K_d$ (nM) |
| --- | --- | --- | --- | --- | --- | --- |
| OsExo70B1 | 10 | AVR-Pii | 100 | 25 | n.b. | n.b |
| OsExo70F2 | 10 | AVR-Pii | 100 | 25 | $-24.04 \pm 0.024$ | $< 1$ |
| OsExo70F3 | 10 | AVR-Pii | 100 | 25 | $-17.67 \pm 0.007$ | 4.19 |

**Table S2.**
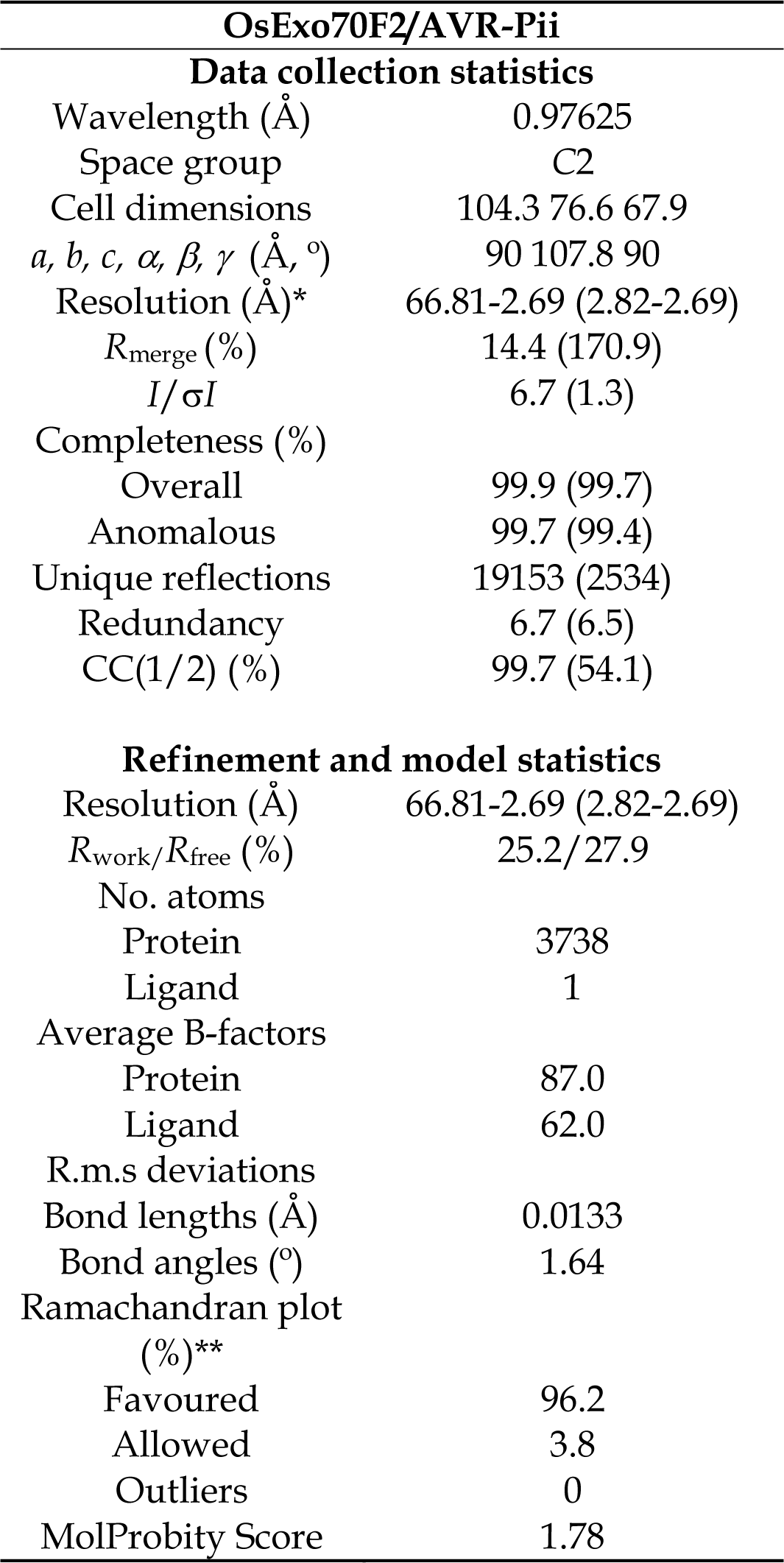
Data collection and refinement statistics. (*) The highest resolution shell is shown in parenthesis. (**) As calculated by MolProbity.

**Table S3.**
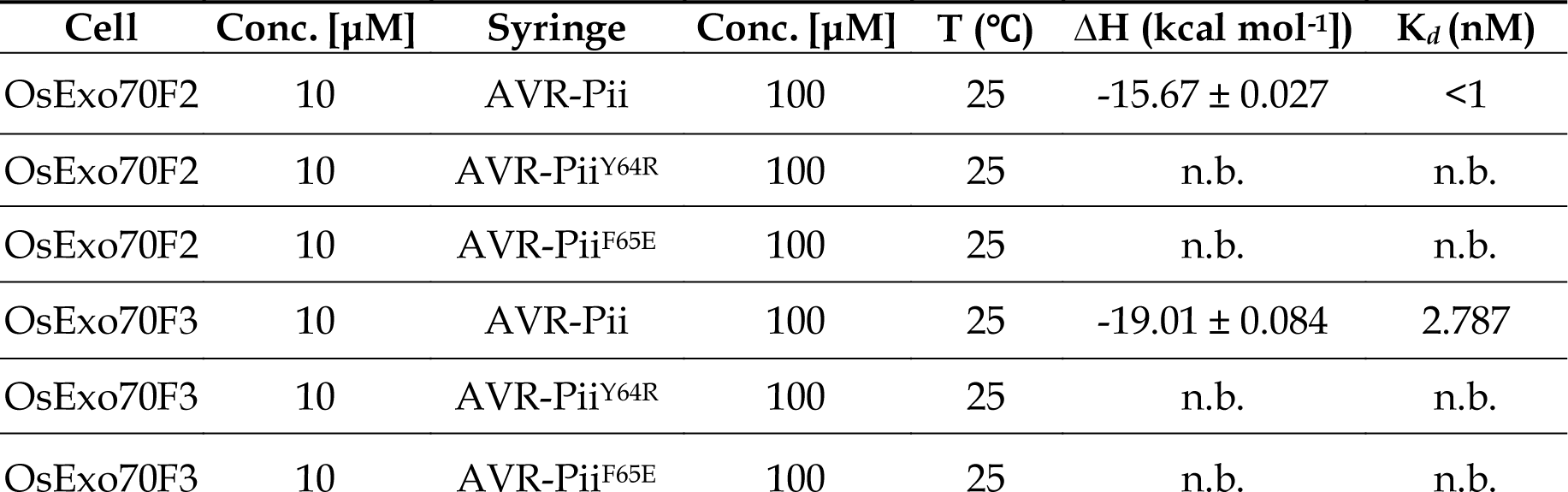
Experimental details and thermodynamic parameters obtained from ITC experiments presented in **Figure 3**.

